# Widespread Polycistronic-like Transcription of the *Populus* Nuclear Genome

**DOI:** 10.1101/2025.07.17.665350

**Authors:** Reed Arneson, William Wittstock, Emma Burke, Aimee Marceau, Yinan Yuan

## Abstract

Transcripts spanning multiple gene loci are common in prokaryotes as a feature of polycistronic gene expression but have traditionally been considered rare in eukaryotic nuclear genomes. In this study, using Nanopore direct RNA sequencing (DRS), we identified widespread mRNA transcripts spanning two or more nuclear gene loci in two *Populus* species, *Populus trichocarpa* (Nisqually-1) and the hybrid poplar ‘717’ (*P. tremula* × *P. alba*). These novel multi-gene-spanning transcripts structurally resemble polycistronic RNAs and are predicted to encode open reading frames (ORFs), including modified and fusion ORFs. Many of these transcripts exhibit tissue-specific, allele-specific, or drought-responsive expression pattern, suggesting potential roles in plant development and environmental adaptation. Functional enrichment and protein localization analyses revealed that genes associated with organelles and membranes are significantly overrepresented within these polycistronic-like (PC-like) transcriptional units. PC-like RNAs also undergo extensive alternative splicing and possess longer polyadenine [poly(A)] tails and fewer N⁶-methyladenosine (m⁶A) modification sites than their monocistronic counterparts. Under drought, dicistronic-like RNAs were more strongly coregulated with the corresponding 5′ monocistronic gene than with the 3′ monocistronic gene, whereas the two corresponding monocistronic genes exhibited a weaker positive correlation in expression. These findings indicate that transcription and RNA processing at PC-like loci are regulated through mechanisms distinct from those governing prokaryotic polycistronic operons. Collectively, our results reveal pervasive PC-like transcription in plant nuclear genomes and highlight its distinctive molecular features, complex regulation, and potential roles in plant growth and environmental adaptation.

## INTRODUCTION

Prokaryotic RNAs are typically transcribed from a single transcription unit containing multiple adjacent and functionally related genes within one RNA molecule encoding multiple proteins, a process known as polycistronic transcription [1, 2]. This arrangement enables genes involved in the same pathway to be co-regulated and co-expressed. Such coordinated expression allows prokaryotes to respond efficiently to environmental and physiological changes and serves as an important mechanism of gene regulation.

Polycistronic-type transcription spanning several neighboring gene loci also occurs in the organelle genomes of eukaryotic cells, as both mitochondria and chloroplasts evolved from prokaryotic ancestors [3, 4]. In contrast, polycistronic-type transcription has traditionally been considered rare in eukaryotic nuclear genomes, where genes are typically transcribed in a monocistronic manner, with each mRNA corresponding to a single gene and encoding a single protein. However, over the past few decades, polycistronic gene loci within nuclear genomes have been identified in several eukaryotes [5–8]. More recently, with advances in long-read transcriptome sequencing, a growing number of polycistronic RNAs overlapping two or more ORFs have been discovered in fungi [9], green algae [10], and plants [11, 12]. These findings suggest that polycistronic-type transcription in eukaryotic nuclear genomes may be far more widespread than previously recognized.

Unlike the polycistronic expression observed in prokaryotes, most polycistronic gene loci identified in eukaryotes appear to produce both monocistronic and polycistronic RNAs. As with any newly characterized RNA type, questions have been raised about the true nature of these polycistronic RNAs in eukaryotes, including concerns about potential sequencing artifacts associated with cDNA sequencing or transcriptional readthrough noise. In green algae, however, experimental evidence has confirmed the existence of gene loci that exclusively produce bona fide polycistronic RNAs capable of generating two distinct proteins from a single dicistronic RNA molecule [10]. Despite these findings, the transcriptional complexity of polycistronic loci in eukaryotes remains poorly understood, particularly their structural and functional diversity, likely due to their low expression levels and the limitations of short-read sequencing technologies employed.

Nevertheless, polycistronic transcription adds another layer of complexity to gene expression regulation in eukaryotes, alongside alternative splicing, alternative polyadenylation, and RNA base modifications. A deeper understanding of eukaryotic polycistronic RNAs, including their structure, expression pattern and functional relevance in key biological and cellular processes, will not only address existing concerns but also provide new insights into the broader gene regulatory networks in eukaryotes.

In this report, we present a genome-wide analysis of polycistronic-type transcription in *Populus*, a model tree species, using extensive Nanopore Direct RNA sequencing (DRS) data from drought experiments involving two *Populus* species. Nanopore DRS sequences native RNA molecules directly [13], thereby avoiding the artifacts commonly associated with cDNA-based sequencing platforms. This technology provides full-length transcript sequences while preserving information on RNA base modifications and 3ʹ end poly(A) tail length [14, 15], making it the most accurate and information-rich approach currently available for polycistronic RNA analysis. Our study identified thousands of polycistronic-type transcriptional units in the *Populus* nuclear genome, revealing their diverse structural and chemical characteristics, dynamic transcriptional patterns, and functional relevance in plant growth and drought stress responses.

## RESULTS

### Identification of widespread polycistronic-type transcription in *Populus*

We included two common *Populus* species in our studies. *P. trichocarpa* was the first tree species to have its genome sequenced [16], and served as the reference genome for *Populus* genomic research. The interspecific hybrid ‘717’ is widely used in transgenic studies due to its well-established in vitro transformation system, and its genome sequence consisting of two haplotype sub-genomes has also recently become available [17]. The effectiveness of drought treatments employed in this study has been previously verified with Illumina short-read sequence analysis (Supplemental Material).

Nanopore DRS reads generated from both Nisqually-1 and 717 were mapped to their respective reference genomes (mapping statistics and run information can be found in Supplemental Table S1), and the resulting alignments were used to identify multi-gene-spanning PC-like RNAs (METHODS). Reads spanning adjacent genes with high coding-sequence homology were excluded unless independent evidence, such as continuous transcription across the intergenic region, supported co-transcription of the corresponding loci. This filtering step was implemented to minimize false-positive PC-like RNA calls arising from minimap2 read-splitting and misalignment artifacts. It also reduced mapping errors associated with allele-specific transcripts of leucine-rich repeat (LRR) genes and other highly similar paralogous sequences, which are particularly prone to incorrect read splitting and misalignment by minimap2.

We first mapped DRS reads generated from Nisqually-1 to the *P. trichocarpa* reference genome. From over 100 million uniquely mapped reads, over 140,000 multi-gene-spanning transcripts were detected, and assigned to 1092 transcription units or loci each supported by multi-gene-spanning reads detected in at least two independent samples. Because these transcripts structurally resemble classical polycistronic transcripts, we define the corresponding loci or transcriptional units as polycistronic-like (PC-like) loci or PC-like transcriptional units. Among these PC-like loci, we identified 578 are shared between root and mature leaf tissue, and 333 are only detected in root and 181 are only present in mature leaf tissue (Supplemental Table S1).

Using a similar procedure, we mapped Nanopore DRS reads from 717 separately to its two haplotype genome assemblies (HAP1 and HAP2) and to the *P. trichocarpa* reference genome. Among approximately 40 million uniquely mapped reads in the HAP1 alignment, more than 112,000 were identified as multi-gene-spanning transcripts, representing 989 PC-like loci (Supplemental Table S1). Notably, approximately 57% of these reads, corresponding to 556 PC-like loci, were also identified as multi-gene-spanning transcripts in the HAP2 alignment (Fig. 1A). Given that the two haplotypes were derived from different *Populus* species, this overlap indicates substantial conservation of gene collinearity and annotated gene structures between the two genomes. At the same time, the remaining haplotype-specific PC-like loci likely reflect allelic sequence variation and structural divergence that contribute to differences in genome annotation between the haplotypes.

**Figure 1.**
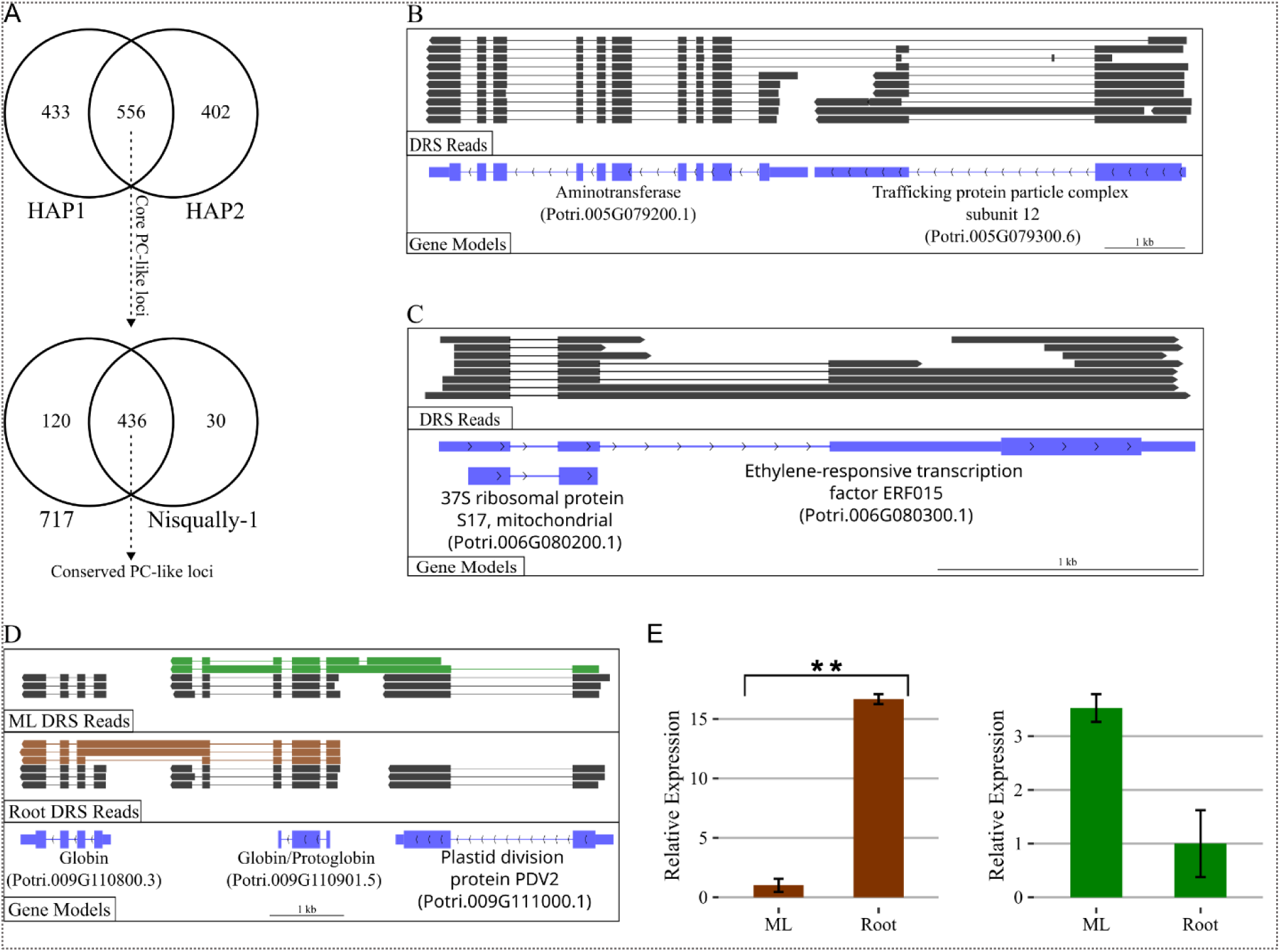
Identification of widespread polycistronic-type transcription in *Populus*. **A.** Venn diagrams showing the numbers of core PC-like loci identified in the two subgenomes of hybrid 717 (HAP1 and HAP2), and the number of loci from this core set shared with *P. trichocarpa* Nisqually-1, representing the conserved PC-like loci between these two species. **B.** An example of a PC-like transcription unit spanning two genes, illustrated by DRS read alignments with defined exon (black boxes) and intron (black lines) structures aligned to the annotated gene models. **C.** An example of an embedded DC-like locus aligned with monocistronic and PC-like DRS reads. **D.** Tissue-specific transcription of overlapping DC-like loci spanning three monocistronic genes and linking distinct monocistronic transcripts in a tissue-specific manner. **E.** Verification of tissue-specific DC-like RNAs transcribed from the overlapping DC-like loci shown in **D** by qPCR analysis. Statistical significance for qPCR data was assessed using an unpaired, two-tailed Student’s t-test assuming equal variances. Data are presented as mean log₂ fold change ± SEM (n = 3 biological replicates). Symbols denote statistical significance: \**p* < 0.05, \*\**p* < 0.01, \*\*\**p* < 0.001.

To ensure a high-confidence dataset, we restricted subsequent analyses to the core set of PC-like loci conserved in both haplotype genomes. Furthermore, more than 400 PC-like loci from this core set were also identified in *P. trichocarpa* genome alignment (Fig. 1A), suggesting that these loci are evolutionarily conserved and may play important biological roles across *Populus* species

One immediate observation regarding the PC-like loci identified in our study is that they are also aligned with pervasive monocistronic transcripts whose boundaries largely matched the annotated gene models of monocistronic counterparts (Fig. 1B), supporting the overall accuracy of the *Populus* genome annotation. This finding is also consistent with previous reports that eukaryotic polycistronic loci frequently generate both monocistronic and polycistronic transcripts. Furthermore, it supports the conclusion that the PC-like RNAs identified in our study are genuine PC-like transcripts extending across gene boundaries, rather than artifacts resulting from inaccurate gene models.

Notably, the PC-like RNAs identified in our study frequently exhibited well-defined exon-intron structures spanning beyond the 3′ untranslated regions (3′ UTRs) of upstream genes and were largely aligned with the gene models of neighboring genes (Fig. 1B). Such processed transcript structures are unlikely to result from random transcriptional readthrough. Instead, they suggest the existence of a previously unrecognized multi-gene-spanning transcriptional mechanism that shares features with polycistronic transcription.

One notable feature of the PC-like transcription units in *Populus* is their characteristic embedded transcriptional architecture, in which PC-like RNAs encompass monocistronic transcripts derived from the same genomic region. This pattern is particularly evident in the embedded dicistronic-like (DC-like) loci identified in our analysis, where one monocistronic gene is nested within a host gene, most commonly within 3′ UTRs, but also identifed in introns and 5’ UTRs. At these loci, the embedded gene, the host gene, or both can also produce monocistronic transcripts alongside DC-like RNAs spanning both genes (Fig. 1C).

Using the *P. trichocarpa* genome as a reference, we classified 114 of the 214 transcribed embedded loci in Nisqually-1 as DC-like loci, based on evidence from both PC-like RNAs and embedded monocistronic transcripts. Although relatively few embedded-gene loci are currently annotated in *Populus* genomes, our results suggest that most are transcribed in a PC-like manner. Together with the general observation of embedded transcript architecture identified in PC-like loci, these findings indicate that embedded-gene organization is a common structural feature of PC-like loci.

The complexity of PC-like transcription units in *Populus* genome is also noteworthy. Although the majority of PC-like RNAs span no more than three adjacent monocistronic gene loci, some complex PC-like transcriptional units are composed of two DC-like loci sharing a monocistronic gene. These overlapping loci give rise to distinct DC-like transcripts encompassing different combinations of adjacent monocistronic genes. As shown in Fig. 1D, genes involved in such overlapping DC-like loci generate distinct DC-like RNAs linking different monocistronic transcripts in leaf and root tissues of Nisqually-1. The tissue-specific expression patterns of these complex transcriptional units were further validated by qPCR analyses (Fig. 1E). These results suggest that transcription at these PC-like loci is likely regulated in a tissue-specific manner and may contribute to tissue-specific biological processes

### Identification of allele-specfic PC-like transcription

Since 717 is an interspecific hybrid and Nisqually-1 is heterozygous in nature [16], it is necessary to probe how allelic variation affects PC-like RNA transcription in *Populus*. Overall, transcription of most PC-like loci, as well as their monocistronic counterparts in Nisqually-1, appeared to be biallelic, with PC-like transcripts commonly detected from both alleles. Nevertheless, allele-specific or allele-biased expression of PC-like RNAs was also frequently observed, particularly in the F1 hybrid poplar 717. To gain a better understanding of such allele-associated transcription, we assigned PC-like reads in 717 to either the *P. tremula* (HAP-1) or *P. alba* (HAP-2) allele based on the number of single-nucleotide polymorphism (SNP) differences detected when the reads were aligned to the two haplotype reference genomes (HAP1, the *P. tremula* subgenome; HAP2, the *P. alba* subgenome) (METHODS).

In 717, analysis of the 556 core PC-like loci revealed that 364 showed biallelic expression, with PC-like transcription detected from both alleles, whereas 188 showed monoallelic expression, with PC-like transcription detected from only one allele. Among the loci with monoallelic expression, 95 showed PC-like transcription from the *P. tremula* allele, whereas 93 showed transcription from the *P. alba* allele. Of the 364 loci with biallelic expression, 123 displayed *P. tremula*-biased PC-like transcription, whereas 150 displayed *P. alba*-biased PC-like transcription, defined as at least a 1.5-fold higher read abundance from one allele than the other.

We also assessed whther these allele-specific PC-like RNAs are drought-responsive by performing differential expression analysis on reads derived from each parental allele. We identified that two monoallele loci, one derived from each parental allele, were differentially regulated under drought conditions (padj < 0.1). In addition, for loci exhibiting biallelic PC-like transcription, we identified 11 biallelic loci with drought-responsive *P. tremula*-specific PC-like transcription and another distinct set of 11 biallelic loci with drought-responsive *P. alba*-specific PC-like RNAs (padj < 0.1) (Supplemental Table S2).

qPCR analysis further confirmed the expression of allele-specific drought-responsive PC-like loci. Among these, a monoallelic PC-like transcription unit originating from the *P. alba* allele was of particular interest. This locus encompasses a gene involved in plastid movement at the 5′ end and a sugar transporter gene at the 3′ end (fig. 2A). The PC-like transcripts were expressed exclusively from the *P. alba* (HAP2) allele and exhibited drought-responsive induction (Fig. 2B).

**Figure 2.**
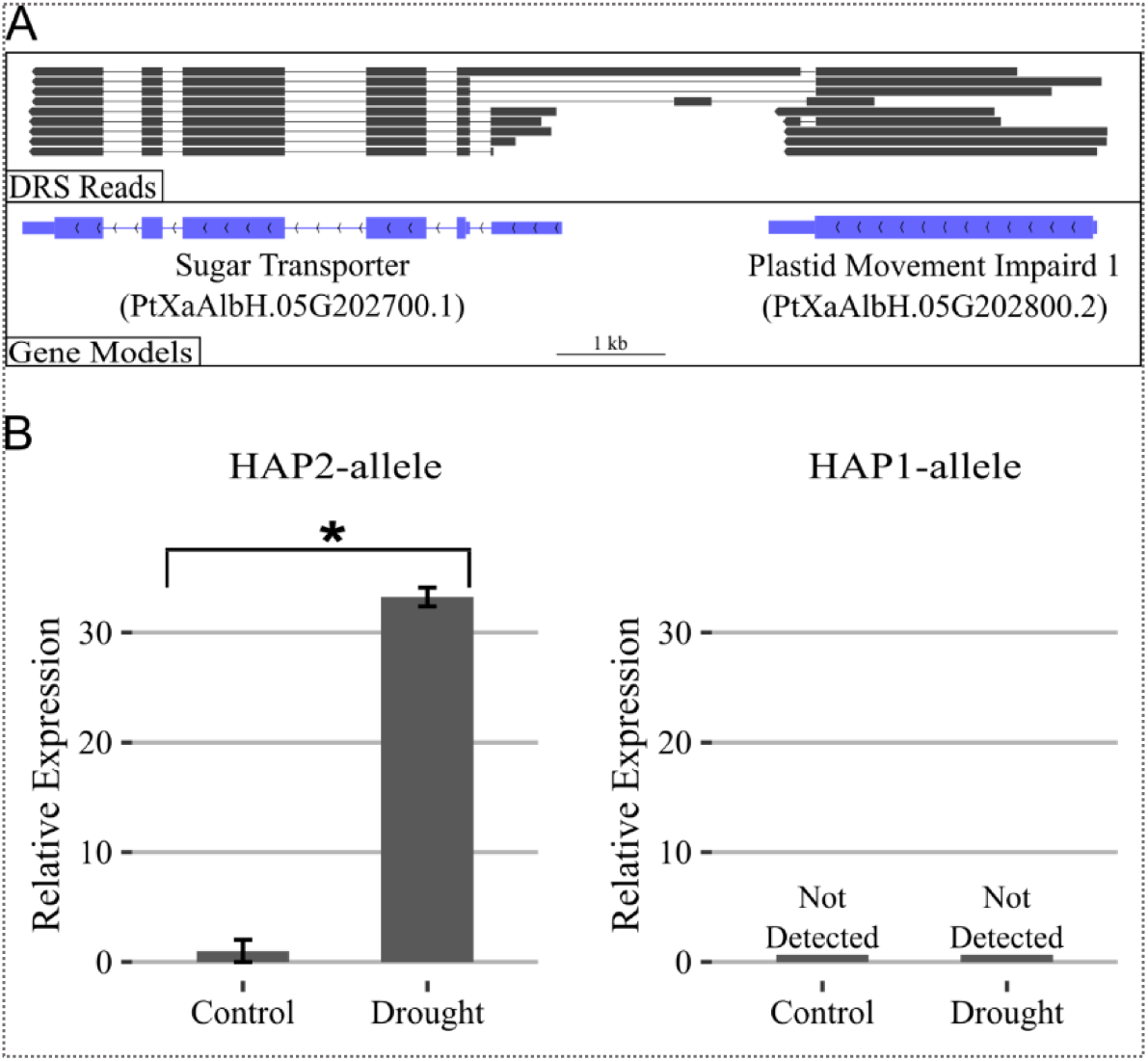
**Allele-specific transcription of DC-like RNAs in the hybrid poplar 717**. **A**. An example of a HAP2 allele-specific DC-like locus involving the sugar transporter and plastid movement genes. **B.** Verification of the drought responsiveness of the HAP2-allele DC-like RNAs shown in **A** by qPCR analysis. Statistical significance for qPCR data was assessed using an unpaired, two-tailed Student’s t-test assuming equal variances. Data are presented as mean log₂ fold change ± SEM (n = 3 biological replicates). Symbols denote statistical significance: \**p* < 0.05, \*\**p* < 0.01, \*\*\**p* < 0.001.

Although both the plastid movement gene and the sugar transporter gene have been implicated in plant stress responses [18–20], our finding provides the first evidence for a potential coregulatory link between sugar transport and chloroplast movement at transcriptional level. Investigating this novel association will advance our understanding of the coordinated regulation of chloroplast dynamics during plant responses to drought stress.

Overall, the results of our allele-specific transcription analysis further support the conclusion that the PC-like RNAs identified in this study are not simply products of transcriptional noise. Instead, they represent complex, tightly regulated transcripts with potential functional roles in plant growth, development, and responses to environmental stress.

### Enrichment of PC-like transcription in organelle-associated nuclear genes

Because polycistronic transcription has traditionally been associated with prokaryotic genes, the widespread PC-like transcription identified in our study prompted us to ask whether this mode of transcription is enriched among genes of prokaryotic origin. Both chloroplasts and mitochondria, which are derived from ancestral prokaryotes, have continuously transferred fragments of their genomes to the nuclear genome, and some of these transferred genes have become nuclear-encoded while remaining functional in the organelles [21–23]. Given the coevolutionary nature of this process, such genes may have retained polycistronic transcription if this feature confers a selective advantage and contributes to plant survival.

To assess whether nuclear-encoded, organelle-associated genes are enriched among genes exhibiting PC-like transcription, we first used a machine-learning tool [24] to predict the subcellular localization of proteins encoded by genes encompassed by PC-like RNAs. Across datasets, approximately 21% of monocistronic genes associated with PC-like transcription were predicted to encode proteins containing organelle-targeting signals, compared with approximately 16% of genes in a control set lacking evidence of polycistronic-type transcription (Fig. 3A). These results indicate that organelle-targeted genes are more likely to be associated with PC-like transcription. Although genes predicted to contain nuclear-targeting signals also appeared to be enriched among PC-like transcripts relative to the control set, prediction of nuclear localization signals is generally less robust than prediction of organelle-targeting signals. Therefore, this observation was not considered further.

**Figure 3.**
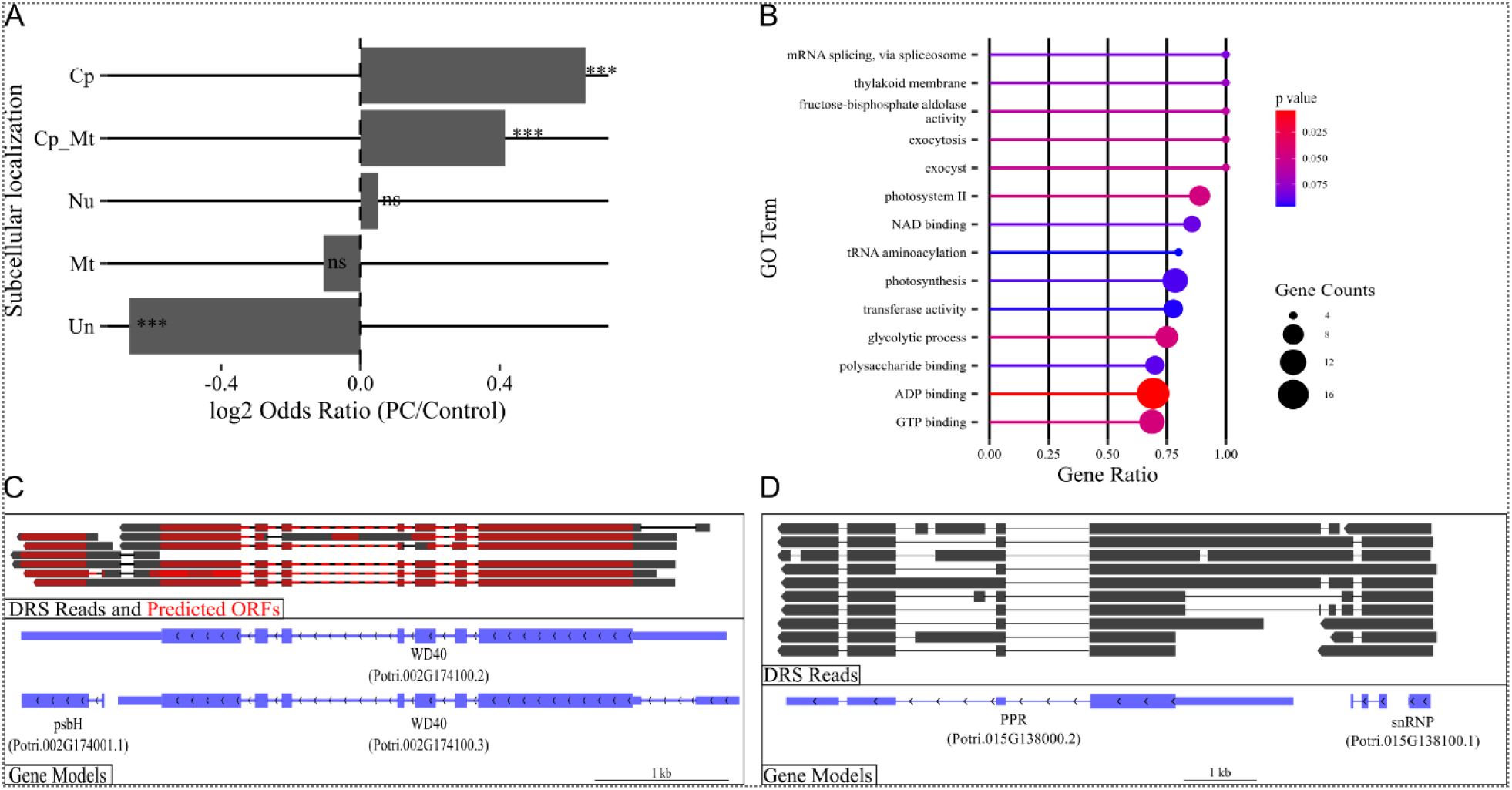
**Subcellular localization prediction and function annotation analysis of PC-like RNAs**. **A**. Predicted subcellular localization of genes spanned by PC-like RNAs, based on encoded signal peptides targeting cellular organelles: Cp, chloroplast; Mt, mitochondria; Cp_Mt, chloroplast or mitochondria; Nu, nucleus; Un, unclassified (no targeting signals detected). Percentages of genes in each category are compared with those of control genes lacking detectable polycistronic-type transcription. Significance test. **B.** GO enrichment analysis of genes spanned by PC-like RNAs in the Nisqually-1 mature leaf dataset using GOseq. Enriched GO terms are displayed based on the specified *P*-value threshold and gene ratios. The gene ratio on the x-axis represents the proportion of genes associated with each GO term among all PC-like RNA-associated genes, and the corresponding gene counts for each GO term are also indicated. **C.** DRS read alignments and ORF predictions (red) for PC-like RNAs spanning the NUPT *psbH* locus and a nuclear gene encoding a WD40 repeat protein. **D.** DRS read alignments supporting the co-transcription of a PPR gene (Potri.015G138000) and a small nuclear ribonucleoprotein gene (Potri.015G138100) through PC-like transcription.

Gene Ontology (GO) analysis of monocistronic genes within PC-like loci further reveals the functional characteristics of the PC-like loci identified in this study. Using the Nisqually-1 mature leaf dataset as an example, we found that the most significantly enriched GO terms were predominantly associated with photosynthesis and chloroplast function (Fig. 3B, Supplemental Table S3). Additional enriched categories include enzymes and transporters containing transmembrane domains, membrane trafficking, signal transduction, transcriptional coregulator activity, and protein kinase and phosphatase activity (Supplemental Table S3).

Further examination of a subset of DC-like loci in which at least one monocistronic gene in each pair is annotated as chloroplast-or mitochondria-associated revealed that nuclear genes with diverse cellular functions are linked to organelle-associated genes. We found that over 60 chloroplast-targeted genes and 50 mitochondria-targeted genes form polycistronic loci with neighboring genes involved in a wide range of cellular processes, including transcriptional regulation, ribosomal structure, chromatin organization, cell-cycle control, RNA binding, and pre-mRNA splicing (Supplemental Table S4). Notably, several nuclear genes encoding photosystem II (PSII) components, such as psbP, psb27, and psbR, produce extensive polycistronic RNA isoforms that link them to genes involved in a range of cellular functions. Although many PSII components are encoded by the chloroplast genome, proper expression of nuclear-encoded PSII genes is essential for chloroplast development and efficient photosynthesis [25–28]. Therefore, it is reasonable to propose that the transcription of polycistronic RNAs at these organelle-associated loci may have significant functional consequences for plant growth and development.

Most notably, in *P. trichocarpa* we identified PC-like RNAs spanning a nuclear-integrated plastid-derived gene (NUPT) encoding a protein with 96% amino acid sequence identity to the chloroplast-encoded PSII protein PsbH and a neighboring gene encoding a WD40-repeat protein (Fig. 3C). NUPTs have generally been considered transcriptionally inactive due to the absence of necessary nuclear regulatory elements [29]. However, the polycistronic transcription of the nuclear copy of *psbH* and its adjacent gene suggests a novel mechanism for NUPT expression in plants. Notably, expression of the chloroplast *psbH* gene, which was profiled using a separate DRS library designed to capture organellar RNAs lacking poly(A) tails (Supplemental Methods), was much higher in leaf tissue than in root tissue, consistent with its role in photosynthesis (Supplemental Figs. S3-4). In contrast, expression of the PC-like NUPT *psbH* transcript differed little between root and leaf tissues, suggesting that the NUPT *psbH* gene may have functions distinct from those of the chloroplast *psbH* gene. The potential role of PC-like NUPT *psbH* in photosynthesis or other cellular processes is therefore particularly intriguing and warrants further investigation, which may yield new insights into the functional and evolutionary significance of NUPTs in plant genomes.

Of note, genes associated with RNA splicing, such as small nuclear ribonucleoproteins (snRNPs) and other splicing factors, despite reported functioning primarily in the nucleus, often produce PC-like RNAs that frequently link with genes involved in distinct cellular processes, including organelle-associated genes such as those encoding pentatricopeptide repeat (PPR) proteins. Many PPR proteins play essential roles in post-transcriptional gene regulation within chloroplasts and mitochondria, where they influence RNA editing, splicing, stability, and translation [30, 31]. Our analysis identified PC-like RNAs encompassing both PPR genes and components of the nuclear pre-mRNA splicing machinery, with a representative locus including a PPR gene and an snRNP-related protein shown in Fig. 3D. Together, these findings reveal a previously unidentified link between nuclear and organellar RNA processing and splicing systems.

Taken together, our functional survey indicates that nuclear-encoded, organelle-associated genes, including novel NUPT genes, frequently generate polycistronic-type RNAs and that this transcriptional mode connects diverse functional classes of nuclear genes, potentially contributing to essential biological processes that require coordinated function among organelles, the nucleus, and other cellular compartments within the cytoplasm.

### Alternative splicing and coding potential of PC-like RNA isoforms

PC-like RNAs identified in *Populus* frequently undergo alternative splicing both within monocistronic gene regions and across the intergenic regions between monocistronic genes, resulting in extensive isoform diversity. As an initial step toward evaluating the coding potential of these transcripts, we employed a computational approach using an open reading frame (ORF) prediction tool [32].

Based on the computation analysis (Method), we found PC-like RNAs either encoding ORFs per transcript or lacking detectable ORFs. The majority of them were predicted to encode one or more ORFs per transcript, including canonical ORFs matching the gene models of monocistornic counterparts. However, we found most predicted ORFs were either structurally distinct from the annotated monocistronic ORFs or represented previously unannotated ORFs. We have also identified fusion ORFs comprising exons derived from two adjacent genes. These noncanonical ORFs may possess functions that differ from those of their canonical counterparts.

One example is a DC-like loci consisting of Potri.002G149900 at the 5′ end, which encodes an enzyme related to alkaline ceramidases, with Potri.002G149800 at the 3′ end, which encodes a protein containing an ankyrin repeat domain (Fig. 4A). One major PC-like RNA isoform transcribed from this locus encodes two ORFs corresponding to the 5′ and 3′ monocistronic genes. Interestingly, each ORF lacks one exon present in its monocistronic counterpart, resulting in modified ORFs that may produce proteins with subtly altered amino acid sequences. The modified ORF at the 3′ end of this DC-like unit lacks the chloroplast transit peptide encoded by the first exon of Potri.002G149800, potentially producing a protein that is no longer targeted to the chloroplast. Notably, the RNA isoform encoding these modified ORFs was detected predominantly in drought-stressed samples (Fig. 4B), suggesting a potential role for the DC-like RNA or its encoded proteins in the drought response of *Populus*.

**Figure 4.**
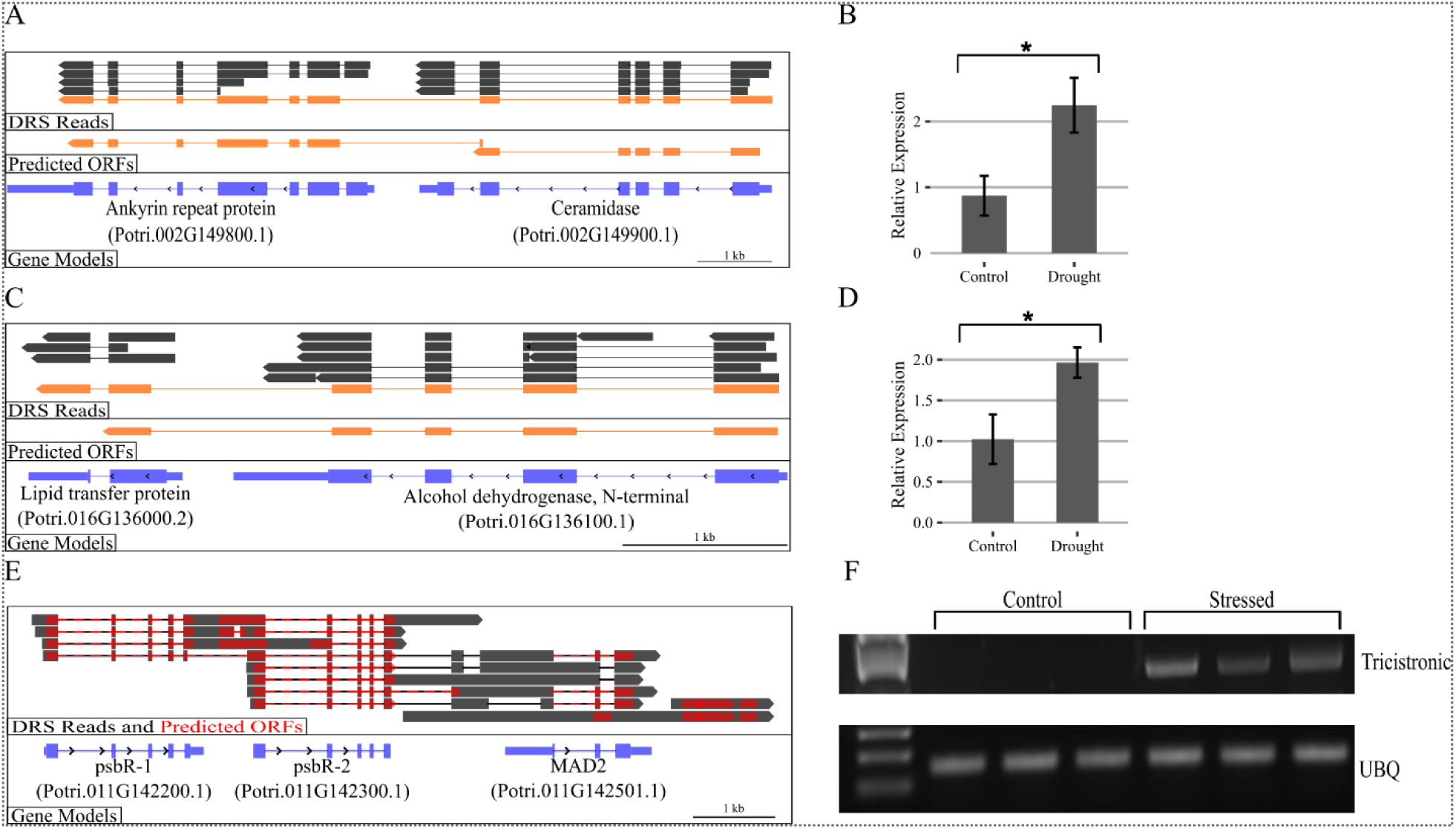
ORF-encoding potential of alternatively spliced PC-like RNA isoforms. **A.** DRS read alignments and the predicted ORF structure of a PC-like RNA isoform encoding two ORFs distinct from the annotated monocistronic ORFs. **B.** qPCR validation of the drought-responsive expression of the PC-like RNA isoform shown in **A**. **C.** DRS read alignments and the predicted fusion ORF structure of a PC-like RNA isoform. **D.** qPCR validation of the drought-induced upregulation of the PC-like RNA shown in **C**. Statistical significance was assessed using an unpaired, two-tailed Student’s t-test assuming equal variances. Data are presented as the mean log₂ fold change ± SEM (n = 3 biological replicates). Statistical significance is indicated as follows: \**p* < 0.05, \*\**p* < 0.01, \*\*\**p* < 0.001. **E.** Alternative splicing of complex PC-like transcription units generates diverse ORF architectures (red), as illustrated by DRS read alignments (gray) spanning monocistronic genes and their intervening intergenic regions. Reads are from mature leaf dataset of the Nisqually-1. **F.** Detection of tricistronic RNAs by semiquantitative RT-PCR on a 1% agarose gel using the ubiquitin gene (UBQ) as an internal control.

The identification of fusion ORFs containing partial or the complete coding sequences of two adjacent monocistronic genes within a single transcript is also noteworthy. One fusion ORF is shown in Fig. 4C, composed of most exons from Potri.016G136100 at the 5′ end, which encodes the N-terminal region of an alcohol dehydrogenase, and Potri.016G136000 at the 3′ end, which encodes a lipid transfer protein (LTP). The predicted length of the fused ORF is 464 amino acids, compared with 384 and 121 amino acids for the ORFs encoded by the 5′-end and 3′-end monocistronic genes, respectively. Both alcohol dehydrogenases and LTPs are well known to be stress-responsive in plants: alcohol dehydrogenases function in oxidation–reduction (redox) reactions [33], while LTPs are involved in lipid metabolism and signaling [34]. Notably, the polycistronic RNA encoding the fusion ORF was significantly more abundant in drought-stressed samples than in controls (Fig. 4D), implying its function in plant response to drought.

When complex PC-like transcription units undergo alternative splicing, diverse isoforms, including DC-like, tricistronic-like (TC-like), and intermediate transcripts, are identified with the potential to produce varied ORF architectures. One such PC-like transcription unit comprises two *psbR* genes, whose products are essential components of the PSII complex [35], and a gene encoding MAD2, a spindle assembly checkpoint protein essential for accurate cell-cycle progression [36] (Fig. 4E). Among the alternatively spliced PC-like RNA isoforms identified at this transcription unit, a TC-like isoform is predicted to encode three ORFs: two corresponding to the annotated gene models Potri.011G142300 (psbR-2) and Potri.011G142501 (MAD2), and a third representing a structurally modified ORF of the Potri.011G142200 (psbR-1). Notably, expression of this TC-like RNA isoform, which spans all three genes fully, is particularly enriched in stressed mature leaf tissue (Fig. 4F). Given the essential role of psbR in plant photosynthesis, the alternatively spliced isoforms of this PC-like transcription unit warrant further functional investigation.

### Polyadenylation and base modification of polycistronic RNAs

Most eukaryotic mRNA possess a poly(A) tail, whose length is associated with translation efficiency, mRNA stability, and subcellular localization [37–39]. We analyzed the poly(A) tail lengths of DC-like RNAs in both *Populus* species and compared them with those of RNAs transcribed from the corresponding monocistronic gene pairs within DC-like loci. We found that DC-like RNAs generally contain significantly longer poly(A) tails than their monocistronic counterparts (Fig. 5A). This pattern is consistent across datasets and species, suggesting that it is an intrinsic feature of polycistronic RNAs in *Populus*.

**Figure 5.**
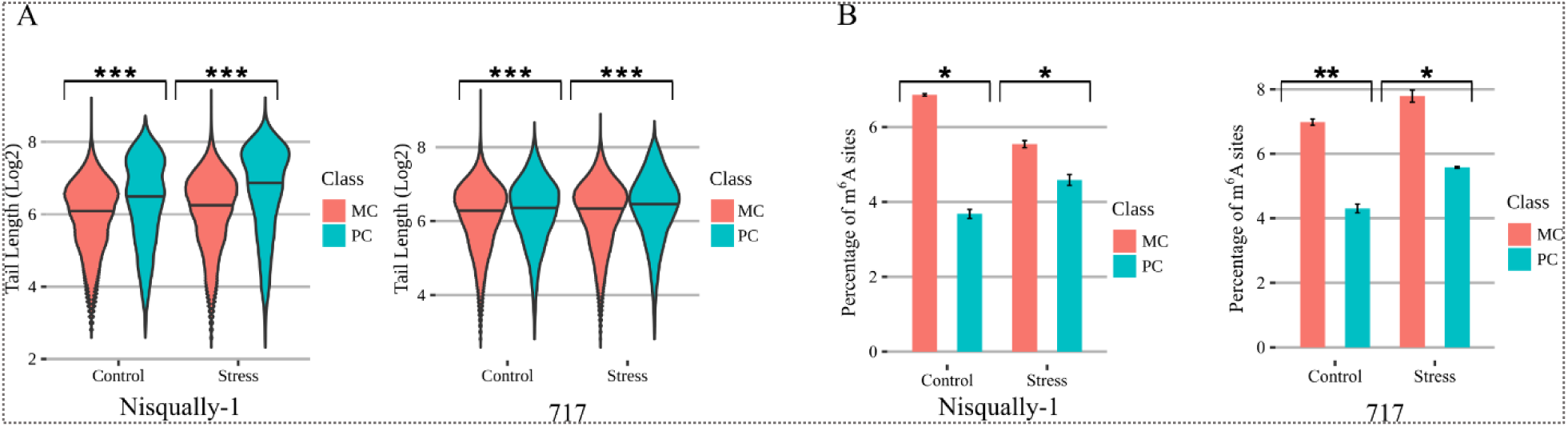
Post-transcriptional characteristics of polycistronic RNAs. **A**. Poly(A) tail length distributions of PC-like RNAs (PC) and monocistronic RNAs (MC) in control and stress samples from the Nisqually-1 root datasets and 717 datasets. Statistical significance was assessed using the Kruskal–Wallis test, followed by pairwise comparisons with Dunn’s test and Bonferroni correction. **B.** Fraction of m⁶A modification sites within the DRACH motif in PC-like (PC) RNAs versus monocistronic (MC) RNAs in both Nisqually-1 (root) and 717 datasets. Statistical significance was determined using an unpaired Student’s t-test assuming equal variance. Symbols denote statistical significance: \**p* < 0.05, \*\**p* < 0.01, \*\*\**p* < 0.001.

In addition, poly(A) tail length appeared to be responsive to drought stress: both monocistronic and PC-like RNAs showed increased tail lengths under drought, with a more pronounced increase observed in PC-like RNAs identified in the Nisqually-1 dataset (Fig. 5A).

We also examined RNA base modifications in DC-like RNA reads, taking advantage of the Nanopore DRS data to directly detect such modifications. Among known RNA modifications, m⁶A RNA base modification is the most prevalent epigenetic mark associated with RNA metabolism and plays an important role in regulating gene expression and plant development [40, 41]. We evaluated m⁶A levels within DRACH motifs in PC-like RNA reads derived from DC-like loci and compared them with those in their monocistronic RNA counterparts. We observed a consistent trend across both *Populus* species, where polycistronic RNAs exhibited significantly lower levels of m⁶A modification than their monocistronic counterparts (Fig. 5B).

In conclusion, the PC-like RNAs identified in *Populus* exhibit distinct post-transcriptional signatures that differentiate them from their monocistronic counterparts. Because these features are closely linked to RNA metabolism and can be dynamically regulated in response to developmental and environmental cues, our findings suggest that PC-like RNA production is subject to specific cellular regulation and may serve functional roles distinct from those of monocistronic transcripts.

### Regulation of PC-like loci in *Populus*

As noted, the PC-like loci identified in *Populus* are aligned with both monocistronic and PC-like reads. This pattern contrasts with the canonical polycistronic gene expression seen in prokaryotes. Investigating the expression of polycistronic RNAs alongside their monocistronic counterparts may provide insights into the regulatory mechanisms governing these loci.

For these analyses, we focused on gene expression of DC-like loci. We first performed Pearson correlation analyses to determine whether the expression of monocistronic gene pairs within DC-like loci was correlated. The resulting correlation coefficients were compared with those of a control dataset consisting of adjacent genes in the same orientation and separated by intergenic distances comparable to those of DC-like loci, but lacking any detectable polycistronic-type transcription. We observed a positive correlation (r = 0.34 in the Nisqually-1 root dataset), between the expression of monocistronic gene pairs encompassed by DC-like RNAs under drought (adjusted *p*-value ≤ 0.1), which was higher than that observed for the control gene pairs (r = 0.18) (Fig. 6A). This pattern was consistently observed across datasets and species (Supplemental Figure S2A).

**Figure 6.**
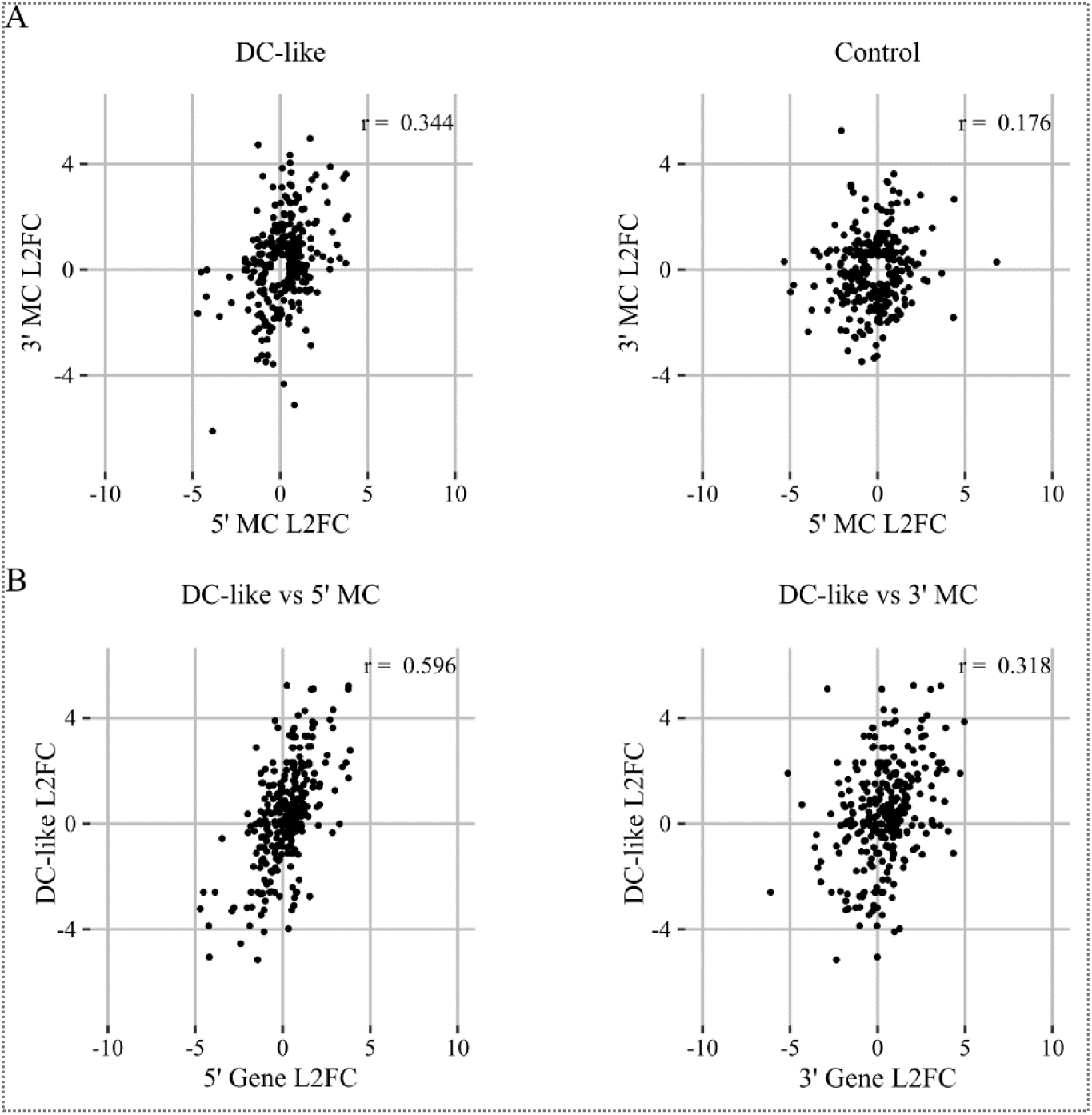
Regulation of PC-like loci in *Populus*. **A.** Pearson correlation analysis of monocistronic (MC) gene pairs within DC-like loci identified in the Nisqually-1 root dataset, compared with a control set of gene pairs from loci with no detectable polycistronic transcription. Only gene pairs in which at least one MC gene was differentially expressed under drought stress (adjusted *p*-value ≤ 0.1) were included. Correlations were calculated using log₂ fold change (L2FC) values. **B.** Pearson correlation analysis between PC-like RNAs and their differentially expressed 5′ MC genes (left) or 3′ MC genes (right) in the Nisqually-1 root dataset (adjusted *p*-value ≤ 0.1). Correlations were calculated using log₂ fold change (L2FC) values.

We next examined the correlation between the expression of DC-like RNAs and that of their corresponding monocistronic counterparts. Specifically, Pearson correlation analyses were performed between DC-like RNA expression and the expression of the corresponding 5′ and 3′ monocistronic genes respectively. Among loci for which at least one monocistronic counterpart was differentially regulated under drought (adjusted *p*-value ≤ 0.1), DC-like RNA expression showed a substantially stronger correlation with the expression of the 5′ monocistronic genes than with that of the 3′ monocistronic genes (r = 0.6 versus r = 0.3 in the Nisqually-1 root dataset) (Fig. 6B). The trend was largely consistent across tissues of Nisqually-1 and between species (Supplemental Figure S2B).

The stronger correlation between the expression of DC-like RNAs and their 5′ monocistronic counterparts provides important insights into the mechanisms underlying polycistronic-like transcription. When considered together with the relatively weak positive correlation observed between the two monocistronic genes within DC-like loci, these findings suggest that expression of the 5′ monocistronic gene plays a key role in the transcription of DC-like RNAs.

Collectively, these observations indicate that the transcription of DC-like RNAs are more closely associated with expression of the 5′ monocistronic gene than with expression of the 3′ monocistronic gene. Future studies should focus on elucidating the molecular mechanisms underlying the coregulation between DC-like RNAs and their 5′ monocistronic counterparts, and their roles in regulating 3’ monocistronic gene expression.

## DISCUSSION

Polycistronic-type transcription in eukaryotic nuclear gene expression has not been systematically characterized. This study fills this gap and provides new insights into its origins, functional mechanisms, and evolutionary significance in coordinating gene expression between plant nuclear and organellar genomes, particularly in the context of plant stress responses.

### The evolutionary origin and functional significance of PC-like transcription

Although polycistronic RNAs have been identified in several eukaryotes, the biological significance of polycistronic-type transcription of nuclear genes remains poorly understood. In the absence of a clear functional role, such transcripts have often been regarded as transcriptional noise, nonfunctional byproducts, or even sequencing artifacts. This skepticism has been reinforced by their generally low abundance and by the technical challenges associated with detecting multi-gene-spanning transcripts using conventional short-read sequencing technologies.

In our study, we identified organelle-associated genes were significantly enriched among PC-like loci. Given that genes in organellar genomes are organized and transcribed as polycistronic units, this enrichment suggests an evolutionary link between PC-like transcription and the transfer of organellar genes to the nuclear genome. PC-like transcription in the nucleus may therefore represent a retained or derived feature associated with the evolutionary integration of organellar genes into the nuclear genetic system. More broadly, these findings place PC-like transcription within the context of nuclear-organelle co-evolution and the integration of gene expression between the nucleus and cytoplasmic organelles.

This evolutionary framework can be traced back to the endosymbiotic events that gave rise to mitochondria and, subsequently, chloroplasts in ancestral eukaryotic cells [42]. The successful integration of these organelles into the host cell required extensive coordination between organellar and nuclear functions. To support this integration, complex endomembrane systems and membrane contact sites (MCSs) evolved to facilitate communication and molecular exchange between organelles and the nucleus [43, 44]. Genes involved in organellar functions and membrane-related processes would therefore be expected to undergo tightly coordinated regulation to support such intercompartmental communication. Consistent with this expectation, our analyses revealed that genes associated with organellar functions and membranes, including many encoding transmembrane proteins involved in transport, signaling, and intracellular trafficking, are enriched among PC-like loci. These findings suggest that PC-like transcription may contribute to the coordinated regulation of genes required for communication and functional integration between organelles and the nucleus.

The evolution of PC-like transcription may have been facilitated by the large reservoir of organelle-derived sequences accumulated within nuclear genomes. Throughout plant evolution, organellar genomes have repeatedly transferred DNA fragments to the nucleus [21]. Some of these transferred sequences have become fully integrated into the nuclear genome as nuclear-encoded, organelle-associated genes, which we found to be significantly enriched among PC-like loci. Others persist as nuclear organellar DNA sequences, including NUPTs and nuclear mitochondrial DNAs (NUMTs). Although the mechanisms underlying organellar DNA transfer and the long-term persistence of NUPTs and NUMTs remain incompletely understood, these sequences are widely recognized as an important source of genetic innovation for nuclear genome evolution [45].

This evolutionary potential is exemplified by our discovery of a novel PC-like locus in the *P. trichocarpa* genome, in which a nuclear copy of *psbH*, a novel NUPT, is linked to a neighboring nuclear gene through PC-like transcription. This finding raises the possibility that organelle-derived DNA fragments can be incorporated into emerging transcriptional units and may acquire novel regulatory or functional roles through association with neighboring genes. More broadly, our results suggest that polycistronic-type transcription may represent an underappreciated mechanism through which organelle-derived sequences are integrated into nuclear gene networks, providing a potential route for evolutionary innovation in plant growth, development, and stress adaptation.

In conclusion, our findings support a model in which PC-like transcription represents an evolutionarily conserved regulatory mechanism that facilitates coordination between cellular compartments, thereby contributing to plant development, environmental adaptation, and the functional integration of nuclear and organellar genomes.

### Regulation and cellular function of PC-like RNAs

Our study raises important questions regarding the regulatory mechanisms governing PC-like loci, which generate both monocistronic and PC-like RNA transcripts. The complex expression relationships observed among monocistronic gene pairs and between PC-like RNAs and their monocistronic counterparts differ markedly from the coordinated expression patterns typically associated with polycistronic genes in prokaryotes. Elucidating the regulatory mechanisms underlying this transcriptional architecture will therefore be a major challenge.

Nevertheless, our findings provide an important starting point. We found that the expression of DC-like RNAs is strongly correlated with the expression of the corresponding 5′ monocistronic gene, suggesting a close regulatory relationship between them. A key unresolved question is whether monocistronic and PC-like transcripts are driven by a shared promoter or regulated through distinct promoter elements. Genome-wide mapping of promoter-associated chromatin marks, such as trimethylation of histone H3 lysine 4 (H3K4me3), using chromatin immunoprecipitation sequencing (ChIP-seq) could provide valuable insights into the promoter architecture and regulatory activity of PC-like loci. Furthermore, functional interrogation of candidate promoter regions through targeted activation or repression experiments would help elucidate the regulatory networks governing incomplete polycistronic transcription in eukaryotes.

Our study also provides insights into the potential cellular functions of PC-like RNAs. We found that PC-like RNAs in *Populus* possess distinct post-transcriptional features, including longer poly(A) tails and fewer m6A modification sites than their monocistronic counterparts. These differences suggest that PC-like RNAs may undergo distinct modes of RNA processing, stability regulation, or subcellular localization, potentially reflecting cellular functions that differ from those of monocistronic transcripts derived from the same loci. Although genome-wide analysis of the subcellular localization of polycistronic RNAs using proximity labeling (APEX-Seq) [46] remains challenging in plants, biochemical approaches such as cell fractionation [47] combined with Nanopore DRS may help determine their relative subcellular distribution and shed light on their cellular function.

### PC-like transcription is a common feature of the *Populus* nuclear genome

Structural analysis of PC-like transcription units revealed a genomic organization resembling embedded gene loci, with PC-like genes encompasses monocistronic gene units. Embedded gene loci are a common feature of eukaryotic genomes [48] and have been described as independently transcribed genes located within introns or arranged in an antisense orientation [49]. When embedded genes are positioned in the same orientation as their host gene, co-transcription can generate polycistronic transcripts containing multiple coding sequences. A well-characterized example has been reported in vertebrates, where a mitochondrial fatty acid synthesis gene is embedded within the 3′ region of the RPP14 locus and is transcribed as a dicistronic RNA encoding two ORFs [8].

Our analyses confirmed that transcription of the embedded gene loci annotated from the *P. trichocarpa* reference genome frequently occurs in a PC-like manner, resulting in the production of both monocistronic and PC-like RNA isoforms. A representative example is the nuclear *psbH* gene, which is embedded within the 3′ UTR of Potri.002G174100, a gene encoding a WD40-repeat protein. Interestingly, transcripts spanning the embedded *psbH* locus are predominantly polycistronic and are predicted to contain two ORFs corresponding to psbH and the WD40-repeat protein. In contrast, transcripts derived from the coding region of Potri.002G174100 are predominantly monocistronic and contain a single ORF encoding only the WD40-repeat protein (Fig. 3C).

Notably, we have observed that the transcript structures of many annotated monocistronic genes in the *P. trichocarpa* genome resemble that of the embedded psbH locus identified in this study. This pattern is particularly prevalent among genes associated with organellar functions or encoding transmembrane proteins. Collectively, these observations point to a broader and previously unrecognized PC-like gene architecture in the *Populus* genome, highlighting an important avenue for future investigation.

Overall, our data indicate that PC-like transcription units are common in *Populus* nuclear genome and may extend to other plant and eukaryotic genomes. Our study supports the view that eukaryotic nuclear genes exhibit a more polycistronic nature, with greater structural and transcriptional complexity, particularly in regions downstream of the 3′ UTR, than previously recognized. This conclusion is consistent with recent reports suggesting that eukaryotic genomes, including the human genome, display a polycistronic organization due to the widespread occurrence of gene embedding [48]. Recognizing the polycistronic features of plant genes will help decipher the intricate gene regulatory networks underlying plant growth and stress responses and has important implications for future advances in genetic engineering and biotechnological applications.

## METHODS

### Plant materials and drought treatment

Nisqually-1 plants were propagated from stem cuttings requested from Oakridge National Lab, whereas 717 plants were propagated via tissue culture. All poplar plants were grown in 3-gallon pots of professional growing mix (Sun Gro), and maintained at full field capacity (FC) for two months prior to initiating drought treatments in a greenhouse, following an established procedure in our lab [50]. Drought stress was applied by withholding water from the treatment group while continuing to water control plants to maintain full FC. Pots were weighed daily to monitor soil water content.

For 717 plants, drought stress was imposed until the soil water content decreased to ∼40% of FC, which was then maintained for 24 hours. Apex leaf tissues were collected from both drought-stressed and control plants and immediately flash-frozen in liquid nitrogen for RNA extraction.

For Nisqually-1 plants, drought-stressed individuals were maintained at 40% of FC for three weeks. Mature leaf (LPI 7) and root tissue were collected from both stressed and control plants and flash-frozen in liquid nitrogen for RNA extraction.

### Nanopore direct RNA library construction and sequencing

Total RNA was extracted from leaf tissues using the CTAB method [51] and treated with DNase I (Thermo Fisher Scientific, EN0521) to remove residual genomic DNA. RNA concentration and purity were assessed using a Qubit and NanoDrop spectrophotometer, and RNA integrity was verified by standard agarose gel electrophoresis.

Both total RNA and mRNA were used for Nanopore direct RNA library construction (Oxford Nanopore Technologies, SQK-RNA004), following the manufacturer’s instructions. For libraries prepared directly from total RNA, ∼1.5 µg of total RNA was used. For libraries prepared from mRNA enriched using Dynabeads® Oligo(dT)25 (Invitrogen, 61002), ∼300 ng of mRNA was typically used. The yield of direct RNA libraries was generally greater than 40 ng. Sequencing was performed on PromethION RNA flow cells (Oxford Nanopore Technologies, FLO-PRO004RA) using a PromethION 2 Solo (P2 Solo) device connected to a GridION Mk1 as a compute resource and operated with standard MinKNOW software (version 24.11.8). When multiple libraries were sequenced on the same flow cell, the Flow Cell Wash Kit (Oxford Nanopore Technologies, EXP-WSH004) was used between runs to remove residual libraries, following the manufacturer’s protocol with a modification: 1 µl of RNase ONE™ Ribonuclease (Promega, M4261) was added along with 1 µl of the enzyme mix provided in the wash kit.

### Nanopore DRS data acquisition, processing, and mapping to *Populus* genome

Raw pod5 files generated by the P2 Solo sequencer were basecalled using the Dorado software (version 0.9.2+2634e9f) provided by Oxford Nanopore Technologies. The basecalling was performed with the model rna004_130bps_sup@v5.1.0, along with poly(A) tail length estimation and modified basecalling models for m⁶A (m6A_DRACH) using the command: dorado basecaller sup,m6A_DRACH --estimate-poly-a.

Basecalled BAM files were converted to FASTQ using SAMtools fastq command: samtools fastq-T’*’, to retain base-modification tags. The resulting RNA reads were then aligned to the *Populus* reference genome v4 using minimap2 [52] with the parameters:-a-x splice-k14-uf-y. The resulting BAM files were filtered with SAMtools [53] using the command: samtools view-h-F 2324 to exclude unmapped, secondary, and supplementary reads.

### PC-like loci identification and ORF annotation

The filtered BAM files containing reads uniquely aligned to the nuclear genomes were further processed to remove reads that also mapped to the organellar genomes with less than 100 nt of soft clipping, as these likely organellar RNAs. The resulting BAM files were then used to identify PC-like loci using a combination of custom R scripts and standard R packages (https://github.com/Reedgarn/Poly_Cistronic_Detection/). Reads overlapping two or more annotated genes in the same orientation, with a minimum overlap of 100 nt per gene, were classified as PC-like RNAs. Loci were designated as PC-like when such reads were detected in at least two independent libraries. The resulting set of PC-like loci was then subjected to additional filtering to remove loci with high sequence similarity that were spanned by allele-specific RNAs, as these regions are prone to erroneous read splitting and alignment by minimap2, potentially leading to false-positive PC-like transcript calls.

To identify ORFs encoded by polycistronic RNAs, we used an ORF prediction tool [32] (orfipy, https://github.com/urmi-21/orfipy). A minimum ORF length threshold of 17 amino acids was applied.

### Gene expression and correlation analysis

Raw read counts for monocistronic genes were generated using featureCounts [54] with the-L and-s 1 options, against the *P. trichocarpa* v4.1 genome annotation. For polycistronic loci, read counting was performed using outputs from custom-built scripts (https://github.com/Reedgarn). Differential expression analysis for both monocistronic and polycistronic data sets was conducted using DESeq2 [55] with default parameters. Statistical significance was defined as an adjusted *p*-value ≤ 0.1, unless otherwise specified. Heatmaps were generated from DESeq2 normalized count data. These counts were first scaled per gene using the base R package scale() and then plotted using ggplot2.

All Pearson correlation analyses were performed using the corr.test function in R to calculate Pearson’s product-moment correlation coefficients between log₂ fold change values generated by DESeq2. The control dataset used in correlation analysis was generated using a random set of gene pairs with the same orientation but lacking detectable polycistronic transcription, matched to the average genomic distance between monocistronic gene pairs in the identified polycistronic loci.

### Assignment of PC-like RNAs to parental alleles and allele-specific expression analysis

To assign DRS reads from 717 libraries to specific parental alleles, DRS reads were aligned to haplotype genomes (HAP-1 and HAP-2) of ‘717’ respectively using the following command: minimap2-a-x splice - k14-uf –eqx –MD-y. The resulting BAM files were parsed using a custom Python script (https://github.com/Reedgarn) to generate a tabular file containing the name of each read with the length of the read in each HAP genome and the number of mismatches in each genome. The resulting file was loaded into R, where the percent mismatch was calculated for the alignment of every read to each HAP genome by dividing the number of mismatches by the length of the alignment. Assignment to a specific allele required at least one of the HAP alignments for a read to contain a mismatch percentage above 0.9%, at which point the read was classified as the HAP genome with the lower mismatch percentage. In cases of a tie in mismatch percentages or no alignment producing a mismatch percentage above 0.9%, the read was marked as coming from either HAP genome.

The allele-assigned DRS reads were then used for differential gene expression analysis of PC-like genes following the standard DESeq2 procedures in an allele-specific manner.

### Semiquantitative RT-PCR and qRT-PCR analyses

First-strand cDNA was synthesized from 2 µg of DNase I-treated total RNA using Induro® Reverse Transcriptase (New England Biolabs, M0681S). Reverse transcription was carried out in a 20 µL reaction containing 4 µL of 5× RT reaction buffer, 1 µL of 0.5 µg/µL oligo(dT) primers (Thermo Fisher Scientific, 18418012), 1 µL of 10 mM dNTPs (Thermo Fisher Scientific, R0191), and 200 units of reverse transcriptase. The reaction mixture was incubated at 55 °C for 15 min, followed by enzyme inactivation according to the manufacturer’s protocol.

Both semi-quantitative RT-PCRs and real-time qPCRs were performed using gene-specific forward and reverse primers designed for each application (Supplemental Table S5). The *UBQ* gene from *P. trichocarpa* (Potri.011G134200) was used as the reference gene for both assays.

For semi-quantitative PCR, amplification was carried out in a 20 µL reaction containing 10 ng of cDNA template, 1 µL of 10 µM forward primer, 1 µL of 10 µM reverse primer, and 10 µL of DreamTaq PCR Master Mix (2×) (Thermo Fisher Scientific, K1082). The PCR cycling conditions were as follows: an initial denaturation at 95 °C for 2 min, followed by 35 cycles of denaturation at 95 °C for 30 s, annealing at 60 °C for 30 s, and extension at 72 °C for 1 min, with a final extension at 72 °C for 7 min. PCR products were resolved on a 1% agarose gel, stained with GelRed (Biotium, 41003), and visualized under UV light using a GelDoc-It imaging system (UVP).

All qPCR reactions were performed using a StepOne Plus Real-Time PCR System. Reactions were set up in a 10 µL volume containing approximately 8 ng of cDNA template, 0.5 µM of each forward and reverse primer, and 5 µL of PowerTrack™ Master Mix (Thermo Fisher Scientific, A46012). The cycling conditions were as follows: an initial denaturation at 95 °C for 2 min, followed by 40 cycles of 95 °C for 15 s and 60 °C for 60 s, followed by a melting curve analysis. Primers were designed to have a melting temperature of approximately 60 °C and to amplify fragments shorter than 200 bp.

The ΔΔCt method was used to analyze qPCR data. Each condition included two biological replicates, with two technical replicates per biological replicate. Average Ct values of the genes of interest were normalized to the average Ct values of the reference gene UBQ to calculate ΔCt. ΔΔCt values were then obtained by subtracting the average ΔCt of the control samples from the ΔCt of the treated samples. Log₂ fold-change values were calculated and are presented in the bar graph. Error bars represent the standard error calculated from ΔCt values. Statistical significance was assessed using an unpaired Student’s t-test assuming equal variance.

### RNA base modification and poly(A) tail length analysis

m⁶A modifications within DRACH sequence contexts were analyzed using modkit (version 0.5.1) with the following command: modkit summary --filter-threshold A:0.8 --mod-thresholds a:0.98. The fractions of m⁶A sites within the DRACH motif in polycistronic and monocistronic reads mapped to DC-like loci were compared using an unpaired Student’s t-test assuming equal variance. Differences with an adjusted *p*-value < 0.05 were considered statistically significant.

Poly(A) tail length distributions between polycistronic and monocistronic reads aligned to dicistronic loci were compared using the TAILcaller package [56] with default settings. Statistical significance was assessed by TAILcaller using the Kruskal-Wallis test, followed by pairwise comparisons using Dunn’s test with Bonferroni correction for multiple testing. Violin plots for poly(A) tail analysis were generated using ggplot2.

### GO analysis and protein subcellular localization prediction

GO terms were compiled from the *P. trichocarpa* reference genome obtained from Ensembl Plants and from agriGO. GO enrichment analysis was performed using GOseq following the standard GOseq pipeline [57]. GO terms with an adjusted *p*-value < 0.05 were considered significantly enriched. The gene ratio was calculated as the number of genes associated with a given GO term divided by the total number of genes annotated with that term.

Protein subcellular localization and organelle-targeting signals of *Populus* genes were predicted using the machine-learning program LOCALIZER [24]. Proteins predicted to contain targeting signals to the chloroplast, dual targeting signals to the chloroplast/mitochondria or chloroplast/nucleus, or triple targeting signals to the chloroplast/nucleus/mitochondria were classified as chloroplast-associated. Similarly, proteins predicted to contain targeting signals to the mitochondria or dual targeting signals to the mitochondria/nucleus were classified as mitochondria-associated. Proteins predicted to contain targeting signals exclusively to the nucleus were classified as nucleus-associated. Proteins for which no targeting signals were predicted by current algorithms were grouped as unclassified. Bar charts summarizing the localization analysis were generated using ggplot2.

## Data access

Sequencing data generated in this study including raw FASTQ and BAM files have been submitted to NCBI Sequence Read Archive (SRA) under the BioProject accession number PRJNA1354876.

All the custom codes used in this study are available at GitHub (https://github.com/Reedgarn) and can also be accessed via the Supplemental Code file.

## Acknowledgements

We thank Dr. Gerald Tuskan (Oak Ridge National Laboratory) for providing stem cuttings of Nisqually-1. We are also grateful to Dr. Chung-Jui Tsai (University of Georgia) and Dr. Victor Busov (Michigan Technological University) for their valuable feedback on the manuscript. We acknowledge the Linux Group of Information Technology at Michigan Technological University for their assistance with software installation.

This work was supported by research seed grants from Michigan Technological University, the Ecological Science Center, and the College of Forest Resources and Environmental Science (start-up funds to Y.Y.) at Michigan Technological University, as well as the McIntire-Stennis Fund (USDA NIFA project #7008315 to Y.Y.). *Author contributions:* Y.Y.: conceptualization; Y.Y. and R.A.: methodology; R.A., E.B., and A.M.: data collection; Y.Y., R.A., and W.W.: data analysis; Y.Y., R.A., and W.W.: manuscript preparation.

