## Supplemental Code for "Widespread Polycistronic-like Transcription of the *Populus* Nuclear Genome": 1A.Remove_Organelle_Reads.py

```
#In python. Use soft-clipping in the organelle alignment as a cut-off to differentiate genuine organelle alignments to alignments of nuclear integrants of organelle DNA.
import pysam

max_clip = #Set number

paln = pysam.AlignmentFile("Organelle_Aligned.bam", "rb") #r means this is for reading
faln = pysam.AlignmentFile("SC_Organelle_Aligned.bam", "wb", template=paln) #w means this is for writing. The template save the header from the provided alignment object


for read in paln:
  if read.cigartuples is None: #Unaligned reads or any other types without CIGAR are skipped over. The tuple is a list of two numbers separated by a comma. The first is the operation, such as matching, insertion, and soft-clipping (which is 4). The second value is the length of that operation.
    continue

  softclip_start = read.cigartuples[0][1] if read.cigartuples[0][0] == 4 else 0 #Break the tuple up into two elements. If the first is 4 (softclipping), then keep. Otherwise, save as 0
  softclip_end = read.cigartuples[-1][1] if read.cigartuples[-1][0] == 4 else 0 #Same as above, but for last value

  if softclip_start <= max_clip and softclip_end <= max_clip:
    faln.write(read)

paln.close()
faln.close()

quit()
```
