## Supplemental Code for "Widespread Polycistronic-like Transcription of the *Populus* Nuclear Genome": 1B.Remove_Organelle_Reads.sh

```
#Generate a nuclear-exclusive BAM file

#Collect read names of organelle-aligned reads with minimal soft-clipping
samtools view SC_Organelle_Aligned.bam | awk '{print $1}' > Organelle_Names.txt

#Filter the nuclear-aligned BAM with the previous file to generate a read list containing nuclear exclusive reads
samtools view Nuclear_Aligned.bam | awk '{print $1}' | awk 'FNR==NR {exclude [$1]; next} !($1 in exclude)' Organelle_Names.txt - > Non_Organelle_Read_Names.txt

#Filter for nuclear exclusive reads
samtools view -bh -F 4 -N Non_Organelle_Read_Names.txt -q 60 Nuclear_Aligned.bam > Nuclear_Only_SC_Removed.bam
```
