## Supplemental Code for "Widespread Polycistronic-like Transcription of the *Populus* Nuclear Genome": 1A.R_PC_Like_Detection_Same_Orientation.r

```
#THis script will generate PC-Like results. The Source_PC_Det.r contains the function for PC detection
library(tidyverse)
library(rtracklayer)
library(IRanges)
library(GenomicRanges)
library(GenomicAlignments)

#Load in an annotation file and collect the features of interest (in this case it is gene)
anno <- import("Annotation.gff3")
Genes <- anno[anno$type == "gene"]

#Add your BAM file
BAM <- "Nuclear_Only_SC_Removed.bam"

#Load in BAM
GR <- readGAlignments(BAM, use.names=TRUE)

#Load in the program
source("Source_PC_Det.r")

#Run the program on the BAM file. Specify your desired overlap length
df1 <- PC_Det(GR, Genes, min.overlap=100, ignore.nested.genes=TRUE, ignore.strand=FALSE)

#Summarise the data to a count matrix
df2 <- df1 %>%
select(RTT) %>%
group_by(RTT) %>%
summarise(Count = n())

#Save data to a tsv file if desired
write_delim(df1, file="PC_Like_Names.tsv", delim = "\t")
write_delim(df2, file="PC_Like_Summary.tsv", delim = "\t")
```
