## Supplemental Code for "Widespread Polycistronic-like Transcription of the *Populus* Nuclear Genome": 2A.PC_Like_Same_Orientation_Filtering.r

```
#This script details the steps to filtering the PC data. We only considered loci that were present in more than one library. Manual filtering can be performed on the PC data before this step or after this step, if desired. 

#Load libraries
library(tidyverse)

#Load in data
C1 <- read_delim("C1_PC_Like_Summary.tsv", delim = "\t")
C2 <- read_delim("C2_PC_Like_Summary.tsv", delim = "\t")
S1 <- read_delim("S1_PC_Like_Summary.tsv", delim = "\t")
S2 <- read_delim("S2_PC_Like_Summary.tsv", delim = "\t")

#Combine the dataframes and filter for those that are present in two or more libraries
dfF <- data.frame(PC = c(C1$PC, C2$PC, S1$PC, S2$PC)) %>%
group_by(PC) %>%
summarise(Count = n()) %>%
filter(Count >= 2)

#Filter each individual library for only the loci that are present in two or more libraries
C1F <- C1 %>%
filter(PC %in% dfF$PC)
C2F <- C2 %>%
filter(PC %in% dfF$PC)
S1F <- S1 %>%
filter(PC %in% dfF$PC)
S2F <- S2 %>%
filter(PC %in% dfF$PC)

#Save the data
write_delim(C1F, file = "Filtered_BCW_C1_PC_Like_Summary.tsv", delim = "\t")
write_delim(C2F, file = "Filtered_BCW_C2_PC_Like_Summary.tsv", delim = "\t")
write_delim(S1F, file = "Filtered_BCW_S1_PC_Like_Summary.tsv", delim = "\t")
write_delim(S2F, file = "Filtered_BCW_S2_PC_Like_Summary.tsv", delim = "\t")
```
