## Supplemental Code for "Widespread Polycistronic-like Transcription of the *Populus* Nuclear Genome": 3.R_PC_Basic_Statistics_And_DC_Filtering.R

```
#This script shows how to find basic information on the filtered PC data in the same orientation. This includes counting the number of dicistronic loci, tricistronic loci, and other loci numbers. Additionally, this script will produce a dicistronic-exlcusive object that can be saved and analyzed in the future. The examples provided are for those in the same orientation, but the exact same protocol can be used for opposite orientations.

library(tidyverse)

#Count Number of DC/TC/etc from filtered data. Start by loading in the data
C1 <- read_delim(file = "Filtered_BCW_C1_PC_Like_Summary.tsv", delim = "\t")
C2 <- read_delim(file = "Filtered_BCW_C2_PC_Like_Summary.tsv", delim = "\t")
S1 <- read_delim(file = "Filtered_BCW_S1_PC_Like_Summary.tsv", delim = "\t")
S2 <- read_delim(file = "Filtered_BCW_S2_PC_Like_Summary.tsv", delim = "\t")

#Now see the number of cistrons. The str_count() function counts the number of commas, which separates the genes in PC loci from the PC_Det() function. One is added since one comma means 2 genes, two commas mean three genes, and so on.
C1s <- C1 %>%
mutate(Cistron = str_count(PC, ",") + 1) %>%
group_by(Cistron) %>%
reframe(Count = n())
C2s <- C2 %>%
mutate(Cistron = str_count(PC, ",") + 1) %>%
group_by(Cistron) %>%
reframe(Count = n())
S1s <- S1 %>%
mutate(Cistron = str_count(PC, ",") + 1) %>%
group_by(Cistron) %>%
reframe(Count = n())
S2s <- S1 %>%
mutate(Cistron = str_count(PC, ",") + 1) %>%
group_by(Cistron) %>%
reframe(Count = n())

#Filter the df's so only dicistronic loci are present. Set the integer to 2 if you want tricistronic, 3 for 4-cistron, etc.
C1d <- C1 %>%
filter(str_count(PC, ",") == 1)
C2d <- C2 %>%
filter(str_count(PC, ",") == 1)
S1d <- S1 %>%
filter(str_count(PC, ",") == 1)
S2d <- S2 %>%
filter(str_count(PC, ",") == 1)

#Save the data to a file
write_delim(C1d, file = "C1_DC_Like.tsv", delim = "\t")
write_delim(C2d, file = "C2_DC_Like.tsv", delim = "\t")
write_delim(S1d, file = "S1_DC_Like.tsv", delim = "\t")
write_delim(S2d, file = "S2_DC_Like.tsv", delim = "\t")
```
