## Supplemental Code for "Widespread Polycistronic-like Transcription of the *Populus* Nuclear Genome": 4A.Same_Orientation_Gene_Reorder.R

```
#The outputs from PC_Det() include the genes associated with a PC read separated by commas. The order of genes in the comma separated list depends upon the genome annotation, with genes being ordered by their left-most coordinate in the annotation. For those on the positive strand, that produces a list of genes ordered from 5' to 3'. For those on the negative strand, however, this produces a list from 3' to 5'. In order to organize all genes in each PC list in a 5' to 3' direction, you can run this script.
#The examples provided in this script are for DC reads, but it will work for tricistronic, 4-cistron, and greater. This script will not work for PC loci containing genes from opposite orientations.

library(tidyverse)

#Make a function to put 5' and 3' genes in correct orientation. It will flip the order of those on the negative strand (denoted as "-" in our gff3 file)
rearrange_genes <- function(gene_pair, annotation) {
	genes <- strsplit(gene_pair, ",")[[1]]

	strand_info <- annotation$strand[annotation$ID %in% genes]
	
	if(any(strand_info == "-")) {
		genes <- rev(genes)
	}
	return(paste(genes, collapse = ","))
}

#Load annotation file and filter for gene features
anno <- import("Annotation.gff3")
genes <- anno[anno$type == "gene"]
genesdf <- as.data.frame(genes)

#Load in the files to rearrange
C1 <- read_delim("C1_DC_Like.tsv", delim = "\t")
C2 <- read_delim("C2_DC_Like.tsv", delim = "\t")
S1 <- read_delim("S1_DC_Like.tsv", delim = "\t")
S2 <- read_delim("S2_DC_Like.tsv", delim = "\t")

#Now run the function to rearrange
C1$PC <- sapply(C1$PC, rearrange_genes, annotation = genedf)
C2$PC <- sapply(C2$PC, rearrange_genes, annotation = genedf)
S1$PC <- sapply(S1$PC, rearrange_genes, annotation = genedf)
S2$PC <- sapply(S2$PC, rearrange_genes, annotation = genedf)

#Save the data, if desired
write_delim(C1, file = "C1_DC_Like_Oriented.tsv", delim = "\t")
write_delim(C2, file = "C2_DC_Like_Oriented.tsv", delim = "\t")
write_delim(S1, file = "S1_DC_Like_Oriented.tsv", delim = "\t")
write_delim(S2, file = "S1_DC_Like_Oriented.tsv", delim = "\t")
```
