## Supplemental Code for "Widespread Polycistronic-like Transcription of the *Populus* Nuclear Genome": Source_PC_Det.r

```
#This is the general function to discover PC reads from the filtered BAM files. Run the R_PC_Detection_Same_Orientation.r or R_PC_Detection_Opposite_Orientation.r script to get the results of this function.

PC_Det <- function(Alignment, Annotation, min.overlap = 1, ignore.strand = FALSE, ignore.nested.genes = FALSE, save.to.disk = 10000){
  
  on.exit(unlink("./TempFilesForRTT_Func", recursive = TRUE), add = FALSE)
  
  #Create a directory to store temp files to save to disk, decreasing memory consumption
  if (dir.exists("./TempFilesForRTT_Func")) {
  	print("Warning: Directory containing temporary data already exists from a previous use of this program. The files are being removed now.")
    unlink("./TempFilesForRTT_Func", recursive = TRUE)
    dir.create("./TempFilesForRTT_Func")
  } 
  else{
  	dir.create("./TempFilesForRTT_Func")
  }

  #Sort files to reduce help reduce memory/computational power
  Annotation <- sort(Annotation)
  #Alignment <- sort(Alignment)
  
  SeqNames <- names(Alignment)
    
  #Create a list of nested genes for future filtering
  if (ignore.nested.genes) { 
    self_hits <- findOverlaps(Annotation, Annotation, ignore.strand = ignore.strand)
    self_overlapping_genes <- split(queryHits(self_hits), subjectHits(self_hits))
    nested_genes <- Filter(function(x) length(x) > 1, self_overlapping_genes) #filter out anything less than 2 elements long
    nested_genes <- nested_genes[!duplicated(lapply(nested_genes, function(x) sort(unique(x))))] #Above line produces duplicates, so this removes duplicates
  }

  #Compute Overlaps between RNA reads and annotation
  Overlap_Hits <- findOverlaps(Annotation, Alignment, ignore.strand = ignore.strand, minoverlap = min.overlap) #(query = Genes, subject = S64)
  
  #Extract overlaps. Take the gene number and read number and combine into a list where each read number is grouped by which gene it overlaps with
  unique_hits <- split(queryHits(Overlap_Hits), subjectHits(Overlap_Hits)) #qHits represent gene number, while subHits represent read number
  
  #Filter the list so that only those with more than one entry are included
  filtered_list <- unique_hits[lengths(unique_hits) > 1]
  
  if (ignore.nested.genes) {#If ignore.nested.genes is true, filter nested genes
    unique_hits_keys <- sapply(filtered_list, function(x) paste(x, collapse = "-")) #Make a key containing hit-hit-hit... for RTT
    nested_keys <- sapply(nested_genes, function(x) paste(sort(unique(x)), collapse = "-")) #Make a key similar to above for all nested genes
    filtered_hits <- filtered_list[!unique_hits_keys %in% nested_keys] #Filter to remove all nested genes
  }
  
  else { #Save the filtered_list to filtered_hits so that nested.genes = TRUE or FALSE can converge back into one workflow
    filtered_hits <- filtered_list
  }
  
  #Remove uneccesary files and clear garbage
  gc()
  
  # Prepare chunk buffers
  chunk_RTT <- character()
  chunk_Reads <- character()
  chunk_id <- 1

  for (i in seq_along(filtered_hits)) {

      # Lookup values
      overlapping_indices <- filtered_hits[[i]]
      overlapping_features <- Annotation[overlapping_indices]
      
      # Gene IDs for RTT
      RTT_value <- paste(mcols(overlapping_features)$ID, collapse = ",")
      
      # Read name
      read_index <- as.integer(names(filtered_hits)[i])
      Read_value <- SeqNames[read_index]

      # Append to chunk
      chunk_RTT <- c(chunk_RTT, RTT_value)
      chunk_Reads <- c(chunk_Reads, Read_value)

      # Once chunk reaches the limit, write to disk
      if (length(chunk_RTT) >= save.to.disk) {
          chunk_df <- data.frame(
              RTT = chunk_RTT,
              Read_Names = chunk_Reads,
              stringsAsFactors = FALSE
          )
          
          saveRDS(chunk_df, 
                  file = paste0("./TempFilesForRTT_Func/Chunk_", chunk_id, ".rds"))

          # CLEAR MEMORY
          rm(chunk_df)
          chunk_RTT <- character()
          chunk_Reads <- character()
          chunk_id <- chunk_id + 1
      }
  }

  # Save remaining rows (final chunk)
  if (length(chunk_RTT) > 0) {
      chunk_df <- data.frame(
          PC = chunk_RTT,
          Read_Names = chunk_Reads,
          stringsAsFactors = FALSE
      )
      saveRDS(chunk_df, 
              file = paste0("./TempFilesForRTT_Func/Chunk_", chunk_id, ".rds"))
  }
  
  chunk_files <- list.files("./TempFilesForRTT_Func", 
                            pattern = "^Chunk_.*\\.rds$", full.names = TRUE)

  RTT_DF <- do.call(rbind, lapply(chunk_files, readRDS))


  #Return data (stdout) so that it can save in the object declared when function is called
  return(RTT_DF)
}
```
