## Supplemental Code for "Widespread Polycistronic-like Transcription of the *Populus* Nuclear Genome": 1A.DC_Like_Correlation_Between_MC_Genes.R

```
#This file details the correlation analysis of monocistronic reads from genes that are associated with dicistronic loci. It includes the steps to produce the DESeq2 log2FoldChange values for MC and PC data, a control for comparison, and correlation testing.

#Load in libraries
library(readxl)
library(DESeq2)
library(tidyverse)
library(rtracklayer)

#Set theme for plots
theme_set(theme_gray(base_family = "serif"))

#load in featureCount data
df <- read_delim("Feature_Count_Matrix_MC.txt", skip = 1, delim = "\t")

#Prepare a count matrix that is compatible with DESeq2
dfA <- df %>%
column_to_rownames(var = "Geneid") %>%
select(matches(".bam"))
colnames(dfA) <- c("C1", "C2", "S1", "S2")


#Prepare a table for DESe2
colF <- data.frame(Condition = rep(c("Control", "Treated"), each = 2), row.names = colnames(dfA))

#Run DESeq2
dds <- DESeqDataSetFromMatrix(dfA, colData = colF, design = ~Condition)
dds <- DESeq(dds)
res <- results(dds, alpha = 0.05)
summary(res)

#Convert the results object into a df for future work. 
dfRes <- as.data.frame(res) %>%
rownames_to_column(var = "Gene")


#-----------------------------------Prepare PC-Like Information-------------------------------
#Load in the PC-like data
C1 <- read_delim("C1_DC_Like_Oriented.tsv")               
C2 <- read_delim("C2_DC_Like_Oriented.tsv")               
S1 <- read_delim("S1_DC_Like_Oriented.tsv")               
S2 <- read_delim("S2_DC_Like_Oriented.tsv")    

#Combine into one matrix
dfO <- data.frame(PC = unique(c(C1$PC, C2$PC, S1$PC, S2$PC)))
DC_Names <- dfO %>%
separate(PC, into = c("Five_Gene", "Three_Gene"), sep = ",")

#Combine the MC DESeq2 data and PC-like Data
Tempdf <- DC_Names %>%
select(Five_Gene, Three_Gene) %>%
left_join(dfRes, by = join_by("Five_Gene" == "Gene")) %>%
select(Five_Gene, Three_Gene, Five_L2FC = log2FoldChange, Five_padj = padj)
MCdf <- Tempdf %>%
left_join(dfRes, by = join_by("Three_Gene" == "Gene")) %>%
select(Five_Gene, Three_Gene, Five_L2FC, Five_padj, Three_L2FC = log2FoldChange, Three_padj = padj)
#The resulting object is now ready for correlation testing.

#------------------------------------Control------------------------------
#Make a control of gene pairs with no observed DC-like expression and similar intergenic distances to our observed set.

#Generate genomic coordinates, first load in annotation file
anno <- import("Annotation.gff3")
genes <- anno[anno$type == "gene"]
genedf <- as.data.frame(genes)


dfPC <- DC_Names %>%
left_join(genedf, by = join_by("Five_Gene" == "Name")) %>%
select(Five_Gene, Three_Gene, Five_Gene_End = end) %>%
left_join(genedf, by = join_by("Three_Gene" == "Name")) %>%
select(Five_Gene:Five_Gene_End, Three_Gene_Start = start)

#Calculate the average length
AvL <- dfPC %>%
mutate(Average_Length = abs((Three_Gene_Start - Five_Gene_End))/2) %>%
reframe(Med = median(Average_Length))


#Now start working on the control. Start with a list of neighboring gene pairs
gene_names <- genes$Name

paired_genes <- data.frame()

for (i in seq_len(length(genes) - 1)) {
  g1 <- genes[i]
  g2 <- genes[i + 1]
 
  # Check if same chromosome and same strand
  if (as.character(seqnames(g1)) == as.character(seqnames(g2)) &&
      as.character(strand(g1)) == as.character(strand(g2))) {
   
    # Build a row of info
    row <- data.frame(
      gene1 = gene_names[i],
      gene2 = gene_names[i + 1],
      chr = as.character(seqnames(g1)),
      strand = as.character(strand(g1)),
      gene1_start = start(g1),
      gene1_end = end(g1),
      gene2_start = start(g2),
      gene2_end = end(g2)
    )
   
    paired_genes <- bind_rows(paired_genes, row)
  }
}


#Now generate a control, filtering out any that are observed in our DC data. The "Average_Length" column can be changed to help generate a sample set closer to what is observed in the DC data. The sample_n() of should be equall to how many DC are present in the observed data
PGrandTemp <- paired_genes %>%
mutate(Average_Length = abs((gene1_end - gene2_start)/2)) %>%
filter(Average_Length <= 1800 & Average_Length >= 1200) %>% #Filters were manually adjusted to generate similar intergenic lengths
unite("PC", gene1, gene2, sep = ",")

#Apply the function to the control
PGrandTemp$PC <- sapply(PGrandTemp$PC, rearrange_genes, annotation = genedf)

#Filter for the final set
PGrand <- PGrandTemp %>%
filter(!(PC %in% dfO$PC)) %>%
sample_n()#Add Number

#Test the median
median(PGrand$Average_Length)

#Make the df
TempdfC <- PGrand %>%
select(PC) %>%
separate(PC, into = c("Five_Gene", "Three_Gene"), sep = ",") %>%
select(Five_Gene, Three_Gene) %>%
left_join(dfRes, by = join_by("Five_Gene" == "Gene")) %>%
select(Five_Gene, Three_Gene, Five_L2FC = log2FoldChange, Five_padj = padj)
Condf <- TempdfC %>%
left_join(dfRes, by = join_by("Three_Gene" == "Gene")) %>%
select(Five_Gene, Three_Gene, Five_L2FC, Five_padj, Three_L2FC = log2FoldChange, Three_padj = padj)


#-----------------------------Correlation Testing-----------------------------------
#The controls were made previously and the results are copied over.
#Test Correlation
CorMC <- MCdf %>%
filter(! is.na(Five_L2FC), ! is.na(Three_L2FC))

CorCon <- Condf %>%
filter(! is.na(Five_L2FC), ! is.na(Three_L2FC))

#Look at sig
CorMCsig <- CorMC %>%
filter(Five_padj <= 0.1 | Three_padj <= 0.1)

R.Sig.MC <- cor.test(CorMCsig$Five_L2FC, CorMCsig$Three_L2FC)

R.Sig.MC
nrow(CorMCsig) 

CorConsig <- CorCon %>%
filter(Five_padj <= 0.1 | Three_padj <= 0.1)

R.Sig.Con <- cor.test(CorConsig$Five_L2FC, CorConsig$Three_L2FC)

R.Sig.Con
nrow(CorConsig)

ggplot(CorMCsig, mapping = aes(x = Five_L2FC, y = Three_L2FC)) +
geom_point(size = 0.5) +
labs(x = "5' MC L2FC", y = "3' MC L2FC") +
theme(panel.background = element_blank(), panel.grid.major = element_line(color = "grey"), axis.title = element_text(size = 9.8), axis.text = element_text(size = 8.5)) +
annotate("text", x = Inf, y = Inf, label = paste("r = ", round(R.Sig.MC$estimate, 3)), color = "black", size = 8.5/.pt, vjust = 2.3, hjust = 1.2) +
ylim(-7, 6) + 
xlim(-10, 10) + 
coord_cartesian(clip = "off")


ggplot(CorConsig, mapping = aes(x = Five_L2FC, y = Three_L2FC)) +
geom_point(size = 0.5) +
labs(x = "5' MC L2FC", y = "3' MC L2FC") +
theme(panel.background = element_blank(), panel.grid.major = element_line(color = "grey"), axis.title = element_text(size = 9.8), axis.text = element_text(size = 8.5)) +
annotate("text", x = Inf, y = Inf, label = paste("r = ", round(R.Sig.Con$estimate, 3)), color = "black", size = 8.5/.pt, vjust = 2.3, hjust = 1.2) +
ylim(-7, 6) + 
xlim(-10, 10) +
coord_cartesian(clip = "off")
```
