## Supplemental Code for "Widespread Polycistronic-like Transcription of the *Populus* Nuclear Genome": 2A.DC_Like_Correlation_Between_PC_And_MC_Genes.R

```
#This script details the correlation analysis of L2FC values from DC reads and MC reads from featureCounts.

#Load in libraries
library(readxl)
library(DESeq2)
library(tidyverse)
library(rtracklayer)

#Set theme for plots
theme_set(theme_gray(base_family = "serif"))

#-----------------------------------------Prepare PC-Like Information----------------------------------------
#Load in the PC-Like Data
C1 <- read_delim("C1_DC_Like_Oriented.tsv")               
C2 <- read_delim("C2_DC_Like_Oriented.tsv")               
S1 <- read_delim("S1_DC_Like_Oriented.tsv")               
S2 <- read_delim("S2_DC_Like_Oriented.tsv")  

#Combine into one object
dfPC <- C1 %>%
full_join(C2, by = "PC") %>%
full_join(S1, by = "PC") %>%
full_join(S2, by = "PC") %>%
column_to_rownames(var = "PC") 
colnames(dfPC) <- c("C1", "C2", "S1", "S2")

#Replace NA with 0's for the counts
dfPC <- dfPC %>%
replace_na(list(C1 = 0, C2 = 0, S1 = 0, S2 = 0))

#Set up and perform DESEq2
colFP <- data.frame(Condition = rep(c("Control", "Treated"), each = 2), row.names = colnames(dfPC))
ddsPC <- DESeqDataSetFromMatrix(dfPC, colData = colFP, design = ~Condition)
ddsPC <- DESeq(ddsPC)
resPC <- results(ddsPC, alpha = 0.05)
summary(resPC)

#Output to a df
dfresPC <- as.data.frame(resPC) %>%
rownames_to_column(var = "PC")

dfPC2 <- dfresPC %>%
separate(PC, ",", into = c("Five_Gene", "Three_Gene"), remove = FALSE)

dfMP <- dfPC2 %>%
left_join(dfRes, by = join_by("Five_Gene" == "Gene")) %>% 
rename(PC_L2FC = log2FoldChange.x, PC_padj = padj.x, Five_L2FC = log2FoldChange.y, Five_padj = padj.y) %>%
select(PC, Five_Gene, Three_Gene, PC_L2FC, PC_padj, Five_L2FC, Five_padj) %>%
left_join(dfRes, by = join_by("Three_Gene" == "Gene")) %>%
rename(Three_L2FC = log2FoldChange, Three_padj = padj) %>%
select(PC, Five_Gene, Three_Gene, PC_L2FC, PC_padj, Five_L2FC, Five_padj, Three_L2FC, Three_padj)

#-----------------------------Correlation Testing-----------------------------
dfAll <- dfMP 
R.All.Five <- cor.test(dfAll$PC_L2FC, dfAll$Five_L2FC) 
R.All.Three <- cor.test(dfAll$PC_L2FC, dfAll$Three_L2FC)


dfSig <- dfMP %>%
filter(Five_padj <= 0.1 | Three_padj <= 0.1) 

R.Sig.Five <- cor.test(dfSig$PC_L2FC, dfSig$Five_L2FC) 
R.Sig.Three <- cor.test(dfSig$PC_L2FC, dfSig$Three_L2FC) 

ggplot(dfSig, mapping = aes(y = PC_L2FC, x = Five_L2FC)) +
geom_point(size = 0.5) +
labs(y = "Polycistronic L2FC", x = "5' Gene L2FC") +
theme(panel.background = element_blank(), panel.grid.major = element_line(color = "grey"), axis.title = element_text(size = 9.8), axis.text = element_text(size = 8.5)) +
annotate("text", x = Inf, y = Inf, label = paste("r = ", round(R.Sig.Five$estimate, 3)), color = "black", size = 8.5/.pt, vjust = 1.6, hjust = 1.2)+
ylim(-7, 6) + 
xlim(-10, 10) +
coord_cartesian(clip = "off")


ggplot(dfSig, mapping = aes(y = PC_L2FC, x = Three_L2FC)) +
geom_point(size = 0.5) +
labs(y = "Polycistronic L2FC", x = "3' L2FC") +
theme(panel.background = element_blank(), panel.grid.major = element_line(color = "grey"), axis.title = element_text(size = 9.8), axis.text = element_text(size = 8.5)) +
annotate("text", x = Inf, y = Inf, label = paste("r = ", round(R.Sig.Three$estimate, 3)), color = "black", size = 8.5/.pt, vjust = 1.6, hjust = 1.2) +
ylim(-7, 6) + 
xlim(-10, 10) +
coord_cartesian(clip = "off")

ggsave("Updated_BCW_Correlatin_Sig_MCvsPC_3.svg", height = 2.35, width = 2.38, units = "in")
```
