## Supplemental Code for "Widespread Polycistronic-like Transcription of the *Populus* Nuclear Genome": 1A.GO_Terms_DC_Same_Orientation.r

```
#This R script details the GO enrichment analysis of DC-like loci relative to controls with no dicistronic-like expression

#Load libraries
library(GO.db)
library(goseq)
library(readxl)
library(DESeq2)
library(tidyverse)
library(rtracklayer)
library(writexl)

#Load in the filtered and properly oriented DC data
C1 <- read_delim("C1_DC_Like_Oriented.tsv")               
C2 <- read_delim("C2_DC_Like_Oriented.tsv")               
S1 <- read_delim("S1_DC_Like_Oriented.tsv")               
S2 <- read_delim("S2_DC_Like_Oriented.tsv") 

#Combine into one df
dfF <- data.frame(PC = unique(c(C1$PC, C2$PC, S1$PC, S2$PC)))

#Split 5' and 3' genes into separate columns
DCdf <- dfF %>%
separate(PC, into = c("Five_Gene", "Three_Gene"), sep = ",") 

#Make df of genes to be used for GO
DC.GO <- data.frame(Gene = c(DCdf$Five_Gene, DCdf$Three_Gene)) 

#Load in featureCounts results to help build the monocistronic data and provide gene lengths
df <- read_delim("Count_Data.txt", skip = 1, delim = "\t")
dfA <- df %>%
dplyr::select(Geneid, matches(".bam"))
colnames(dfA) <- c("Gene", "C1", "C2", "S1", "S2")
dfCon <- dfA 

#Filter the df to remove DC-like associated genes
Con.GO <- data.frame(Gene = dfCon$Gene) %>%
filter( !(Gene %in% DC.GO$Gene)) %>%
sample_n() #Set # to DC-like genes

#Create a dataframe for goseq, where dicistronic-associated genes are labelled as 1, and monocistronic genes are labelled as 0
Con.GO$Class <- rep("Control", nrow(Con.GO))
DC.GO$Class <- rep("DC", nrow(DC.GO))
GO.Both <- rbind(Con.GO, DC.GO) %>%
mutate(DC = if_else(Class == "DC", 1, 0)) %>%
unique() %>%
arrange(Gene)

#Now produce a length vector for goseq
dfLe <- df %>%
dplyr::select(Geneid, Length) %>%
filter(Geneid %in% GO.Both$Gene) %>%
arrange(Geneid)

#Sanity check
sum(GO.Both$Gene == dfLe$Geneid) == nrow(dfLe)

#make the length vector
vec.length <- as.vector(dfLe$Length)
names(vec.length) <- dfLe$Geneid

#Prepare GO Terms. GO terms can be accessed in multiple locations, including through BiomaRt, agriGO, and Phytozome
GO <- read_delim("GO_Terms.txt", delim = "\t")
GO <- GO %>%
drop_na()
colnames(GO) <- c("gene_id", "go_id")
GO.filt <- GO %>%
filter(gene_id %in% GO.Both$Gene)

#Make vector of Genes
vec.genes <- as.vector(GO.Both$DC)
names(vec.genes) <- GO.Both$Gene

#Run goseq
pwf <-nullp(vec.genes, bias.data = vec.length)
GO.wall.DC <- goseq(pwf, gene2cat = as.data.frame(GO.filt), use_genes_without_cat=TRUE)

#You can save this data to an Excel file 
write_xlsx(GO.wall.DC, path = "GO_Terms_DCvsControl.xlsx")
```
