## Supplemental Code for "Widespread Polycistronic-like Transcription of the *Populus* Nuclear Genome": 1A.Localization_Detection.sh

```
#!/usr/bin/env bash

#This bash script details the use of LOCALIZER.py for protein localization predictions. The primary transcript file from Phytozome was used, but other proteomes can be used as well.

#Run LOCALIZER.py, making sure to specify where the program is located if not added to the PATH variable
python LOCALIZER-master/Scripts/LOCALIZER.py -p -o predictions -i Ptrichocarpa_533_v4.1.protein_primaryTranscriptOnly.fa

#Filter the data to collect just the gene names. Perform for all files predicted
grep ">" predictions/chloroplast_predicted.fasta > Temp_Plastid_Gene_Names.txt

#This command removes the > sign, prints the gene name, and removes additional information appended to the gene name
awk '{print $1}' Temp_Plastid_Gene_Names.txt | sed 's/>//g' - | awk -F '.' '{print $1"."print $2}' - > Chloroplast_Exclusive_Names.txt
```
