## Supplemental Code for "Widespread Polycistronic-like Transcription of the *Populus* Nuclear Genome": 2A.Localization_Of_DC_Like_Associated_Genes.r

```
#This R script details how localization information is combined with the DC-like data.

#Load libraries
library(readxl)
library(DESeq2)
library(tidyverse)
library(rtracklayer)
library(writexl)

#Load in the featureCounts data and perform DESeq2
df <- read_delim("Count_Data.txt", skip = 1, delim = "\t")
dfA <- df %>%
column_to_rownames(var = "Geneid") %>%
select(matches(".bam"))
colnames(dfA) <- c("C1", "C2", "S1", "S2")

#DESeq2
colF <- data.frame(Condition = c("Control", "Control", "Treated", "Treated"), row.names = colnames(dfA))
dds <- DESeqDataSetFromMatrix(dfA, colData = colF, design = ~Condition)
dds <- DESeq(dds)
res <- results(dds, alpha = 0.1)
summary(res)

#Make df out of DESeq2 results
dfRes <- as.data.frame(res) %>%
rownames_to_column(var = "Gene")

#Now load in the DC data to isolate the genes associated with DC loci. For this, we are loading the filtered and ordered DC data
C1 <- read_delim("C1_DC_Like_Oriented.tsv")               
C2 <- read_delim("C2_DC_Like_Oriented.tsv")               
S1 <- read_delim("S1_DC_Like_Oriented.tsv")               
S2 <- read_delim("S2_DC_Like_Oriented.tsv")  

#Now combine the data the loci into one df
dfO <- data.frame(PC = unique(c(C1$PC, C2$PC, S1$PC, S2$PC)))
DC_Names <- dfO %>%
separate(PC, into = c("Five_Gene", "Three_Gene"), sep = ",")

#Combine the MC DESeq2 data and 
Tempdf <- DC_Names %>%
select(Five_Gene, Three_Gene) %>%
left_join(dfRes, by = join_by("Five_Gene" == "Gene")) %>%
select(Five_Gene, Three_Gene, Five_L2FC = log2FoldChange, Five_padj = padj)

MCdf <- Tempdf %>%
left_join(dfRes, by = join_by("Three_Gene" == "Gene")) %>%
select(Five_Gene, Three_Gene, Five_L2FC, Five_padj, Three_L2FC = log2FoldChange, Three_padj = padj)

#Prepare the localization information
CE <- read_delim("Chloroplast_Exclusive_Names.txt", delim = " ", col_names="Gene")
CE$Localization <- rep("Chloroplast", nrow(CE))

CM <- read_delim("Chloroplast_Mito_Names.txt", delim = " ", col_names="Gene")
CM$Localization <- rep("Chloro_Mito", nrow(CM))

CN <- read_delim("Chloroplast_Nuc_Names.txt", delim = " ", col_names="Gene")
CN$Localization <- rep("Chloro_Nuclear", nrow(CN))

ME <- read_delim("Mito_Eclusive_Names.txt", delim = " ", col_names="Gene")
ME$Localization <- rep("Mito", nrow(ME))

MN <- read_delim("Mito_Nuclear_Names.txt", delim = " ", col_names="Gene")
MN$Localization <- rep("Mito_Nuclear", nrow(MN))

NE <- read_delim("Nuclear_Exclusive.txt", delim = " ", col_names="Gene")
NE$Localization <- rep("Nuclear", nrow(NE))

CMN <- read_delim("CMN_Names.txt", delim = " ", col_names="Gene")
CMN$Localization <- rep("CMN", nrow(CMN))

dfL <- rbind(CE, CM, CN, CMN, MN, ME, NE)

#This is missing all of the "no predictions" from LOCALIZER. Combine with the gene list from featureCounts gene list for MC analysis.

TempA <- dfA %>%
rownames_to_column(var = "Gene") %>%
select(Gene)

dfLT <- dfL %>%
full_join(TempA, by = "Gene") %>%
replace_na(list(Gene = "Error", Localization = "Not Classified"))


#Now combine localization with the DC data
DCL <- MCdf %>%
left_join(dfLT, by = join_by("Five_Gene" == "Gene")) %>%
left_join(dfLT, by = join_by("Three_Gene" == "Gene")) %>%
select(Five_Gene:Three_padj, Five_Localization = Localization.x, Three_Localization = Localization.y)

#Create Localization list for all values with no DC expression
DCnames <- unique(c(DCL$Five_Gene, DCL$Three_Gene, dfPCL$Gene))

MCL <- TempA %>%
filter(!(Gene %in% DCnames)) %>%
left_join(dfLT) 


#Summarise the information
dfDCG <- data.frame(Localization = c(DCL$Five_Localization, DCL$Three_Localization))
SumDC <- dfDCG %>%
group_by(Localization) %>%
summarise(Total = n()) %>%
mutate(Class = rep("DC Associated", 8))

#Make into Factor to specify order
Organelle_Order <- c("Chloroplast", "Chloro_Nuclear", "Chloro_Mito", "Mito", "Mito_Nuclear", "Nuclear", "CMN", "Not Classified")
SumDC$Localization <- factor(SumDC$Localization, levels = Organelle_Order)

SumMC <- MCL %>%
select(Localization) %>%
group_by(Localization) %>%
summarise(Total = n()) %>%
mutate(Class = rep("No DC Expression", 8))
SumMC$Localization <- factor(SumMC$Localization, levels = Organelle_Order)

dfA <- rbind(SumDC, SumMC)
write_xlsx(dfA, path = "BCW_Total_All_DC_Localization.xlsx")

#You can use the summary information in Excel to investigate localization, which can then be graphed. You can also use the DESeq2 data to look for drought-specific changes in protein localizations.

#The statistical testing and graphing were performed as follows:
theme_set(theme_gray(base_family = "serif"))


library(dplyr)
library(purrr)
library(ggplot2)
dfB <- read_excel(path = "BCW_Total_All_DC_Localization.xlsx", sheet = "Number_Bar")

df <- dfB

#Generate enrichment tables to and test for significance using a two-tailed fisher test
total_pc <- sum(df$PC_Counts)
total_ct <- sum(df$Cont_Counts)

calc_stats <- function(i) {
  
  a <- df$PC_Counts[i]
  c <- df$Cont_Counts[i]
  
  b <- total_pc - a
  d <- total_ct - c
  
  mat <- matrix(c(a, b,
                  c, d),
                nrow = 2,
                byrow = TRUE)
  
  ft <- fisher.test(mat)
  
  or <- (a / b) / (c / d)
  
  tibble(
    Localization = df$Localization[i],
    log2OR = log2(or),
    p = ft$p.value,
    odds_ratio = ft$estimate
  )
}

res <- map_dfr(seq_len(nrow(df)), calc_stats)

#Correct for multiple test correction using Benjamini-Hochberg
res <- res %>%
  mutate(
    padj = p.adjust(p, method = "BH"),
    sig = case_when(
      padj < 0.001 ~ "***",
      padj < 0.01  ~ "**",
      padj < 0.05  ~ "*",
      TRUE ~ "ns"
    )
  )


ggplot(res, aes(x = reorder(Localization, log2OR), y = log2OR)) +
  geom_col() +
  coord_flip() +
  geom_hline(yintercept = 0, linetype = "dashed") +
  geom_text(aes(label = sig), hjust = -0.2, size = 8.5, size.unit = "pt") +
  labs(
    x = "Subcellular localization",
    y = "log2 Odds Ratio (PC/Control)") +
  theme(axis.title = element_text(size = 9.8), axis.text = element_text(size = 8.5), panel.grid.major.x = element_blank(), panel.grid.minor = element_blank(), panel.grid.major.y = element_line(color = "black"), panel.background = element_blank())
```
