## Supplemental Code for "Widespread Polycistronic-like Transcription of the *Populus* Nuclear Genome": 1A.PC_Tail_Length_File_Prep.sh

```
#!/usr/bin/env bash

#This script details the steps to generate a files to compare the tail length of genes associated with PC loci and those with no PC detection.
#A file containing tail lengths and read names is needed. This file should contain two columns, with the read names in the first column and the tail length in the second, separated by a space or tab.
#The CORE file formats for nuclear aligned reads are also needed if you want to produce a set of monocistronic data from polycistronic-associated genes. Ensure that the featureCounts data is strand specific and does not count reads that align to multiple features.

#Collect FC Assigned reads
grep 'Assigned' Nuclear_Only_SC_Removed.bam.featureCounts > MC_Names.txt


#Use the PC-like names from PC_Det program to make PC read names.
awk '{print $2}' Filtered_BCW_C1_PC_Like_Names.tsv > PC_Names.txt


#Now add the tail length to each file type.
awk 'NR==FNR {keep[$1]; next} $1 in keep' MC_Names.txt Tail_Lengths.txt > MC_Tail_Lengths.tsv
awk 'NR==FNR {keep[$1]; next} $1 in keep' PC_Names.txt Tail_Lengths.txt > PC_Tail_Lengths.tsv


#Prepare Files for TAILCALLER. First Collect read names and then filter BAM files
awk '{print $1}' MC_Tail_Lengths.tsv > MC_Names.txt
awk '{print $1}' PC_Tail_Lengths.tsv > PC_Names.txt


samtools view -bh -N MC_Names.txt Nuclear_Only_SC_Removed.bam > MC_BAM.bam
samtools view -bh -N PC_Names.txt Nuclear_Only_SC_Removed.bam > PC_BAM.bam
```
