## Supplemental Code for "Widespread Polycistronic-like Transcription of the *Populus* Nuclear Genome": 2A.PC_Tail_Length_Analysis.r

```
#This R script details the comparisons of tail lengths from PC reads and MC reads. The files needed for this are generated in 1A.PC_Tail_Length_File_Prep.sh.

#Work to perform tail length analysis of the filtered read length data
library(tidyverse)
library(Rsamtools)
library(dplyr)
library(stats)
library(tidyr)
library(rlang)
library(ggplot2)
library(ComplexHeatmap)
library(ggtree)
library(circlize)
library(car)
library(dunn.test)
library(nortest)
library(rstatix)
library(stringr)

theme_set(theme_gray(base_family = "serif"))


#Now load in PC data
C1PC <- read_delim("C1_PC_Tail_Lengths.tsv", col_names = c("Name", "Count"), col_select = Count)
C1PC$Library <- rep("C1", nrow(C1PC))
C1PC$Class <- rep("PC", nrow(C1PC))
C1PC$Condition <- rep("Control", nrow(C1PC))

C2PC <- read_delim("C2_PC_Tail_Lengths.tsv", col_names = c("Name", "Count"), col_select = Count)
C2PC$Library <- rep("C2", nrow(C2PC))
C2PC$Class <- rep("PC", nrow(C2PC))
C2PC$Condition <- rep("Control", nrow(C2PC))

S1PC <- read_delim("S1_PC_Tail_Lengths.tsv", col_names = c("Name", "Count"), col_select = Count)
S1PC$Library <- rep("S1", nrow(S1PC))
S1PC$Class <- rep("PC", nrow(S1PC))
S1PC$Condition <- rep("Stress", nrow(S1PC))

S2PC <- read_delim("S2_PC_Tail_Lengths.tsv", col_names = c("Name", "Count"), col_select = Count)
S2PC$Library <- rep("S2", nrow(S2PC))
S2PC$Class <- rep("PC", nrow(S2PC))
S2PC$Condition <- rep("Stress", nrow(S2PC))


#Now load in MC Data
C1MC <- read_delim("C1_MC_Tail_Lengths.tsv", col_names = c("Name", "Count"), col_select = Count)
C1MC$Library <- rep("C1", nrow(C1MC))
C1MC$Class <- rep("MC", nrow(C1MC))
C1MC$Condition <- rep("Control", nrow(C1MC))

C2MC <- read_delim("C2_MC_Tail_Lengths.tsv", col_names = c("Name", "Count"), col_select = Count)
C2MC$Library <- rep("C2", nrow(C2MC))
C2MC$Class <- rep("MC", nrow(C2MC))
C2MC$Condition <- rep("Control", nrow(C2MC))

S1MC <- read_delim("S1_MC_Tail_Lengths.tsv", col_names = c("Name", "Count"), col_select = Count)
S1MC$Library <- rep("S1", nrow(S1MC))
S1MC$Class <- rep("MC", nrow(S1MC))
S1MC$Condition <- rep("Stress", nrow(S1MC))

S2MC <- read_delim("S2_MC_Tail_Lengths.tsv", col_names = c("Name", "Count"), col_select = Count)
S2MC$Library <- rep("S2", nrow(S2MC))
S2MC$Class <- rep("MC", nrow(S2MC))
S2MC$Condition <- rep("Stress", nrow(S2MC))


#Combine the data
dfA <- rbind(C1PC, C2PC, S1PC, S2PC, C1MC, C2MC, S1MC, S2MC)

#Remove extra objects
rm(list = c("C1PC", "C2PC", "S1PC", "S2PC", "C1MC", "C2MC", "S1MC", "S2MC"))

#Log2 transform the tail lengths
dfA <- dfA %>%
mutate(Log2 = log2(Count))

#Plot and save.
ggplot(dfA, mapping = aes(x = Condition, y = Log2, fill = Class)) +
geom_violin(quantiles = c(0.5), quantile.linetype = 1) +
labs(x = NULL, y = "Tail Length (Log2)") +
theme(panel.grid.major.x = element_blank(),
	 panel.background = element_blank(),
	 panel.grid.major.y = element_line(color = "grey"),
	 axis.text = element_text(size = 8.5), axis.title = element_text(size = 9.8),
	 legend.text = element_text(size = 8.5), legend.title = element_text(size = 9.8))
	 
ggsave("PC_BCW.svg", width = 3.065, height = 2.280, units = "in")

rm(list = ls())


#TAILCALLER
bf1 <- "C1_PC_BAM.bam"
bf2 <- "C2_PC_BAM.bam"
bf3 <- "S1_PC_BAM.bam"
bf4 <- "S2_PC_BAM.bam"
bf5 <- "C1_MC_BAM.bam"
bf6 <- "C2_MC_BAM.bam"
bf7 <- "S1_MC_BAM.bam"
bf8 <- "S2_MC_BAM.bam"

samples_table <- data.frame(bam_path = c(bf1,bf2,bf3,bf4,bf5,bf6,bf7,bf8),
                            sample_name = c("C1_PC","C2_PC","S1_PC","S2_PC","C1_MC","C2_MC","S1_MC","S2_MC"), 
			    group = c(rep(c("Control_PC", "Stress_PC"), each = 2), rep(c("Control_MC", "Stress_MC"), each = 2)))

get_polyA_out <- TAILcaller::get_polyA(samples_table = samples_table)
density_plot <- TAILcaller::plot_density(polyA_table = get_polyA_out,stats = "mean",grouping_factor = "group")
Test <- TAILcaller::calculate_statistics(polyA_table = get_polyA_out, grouping_factor = "group", control_group= c("Control_PC", "Control_MC"), treated_group= c("Stress_PC", "Stress_MC"))

#Part of the TAILcaller output are the comparisons between different groups with multiple testing correction. These were used to determine p-values 
```
