## Supplemental Material for "Widespread Polycistronic-like Transcription of the *Populus* Nuclear Genome"

**Figure S1.** Principal component analysis of gene expression in Nisqually-1 and 717 under drought stress, using Illumina short-read RNA-seq data.

**Figure S2.** Correlation analysis of the expression of DC-like RNAs in Nisqually-1 mature leaf and 717

**Figure S3.** DRS read alignments supporting the transcription of the chloroplast genome-encoded *psbH* in leaf and root tissue of *P. trichocarpa*.

**Figure S4.** Bar chart comparing the expression of the NUPT *psbH* gene and the chloroplast *psbH* gene in mature leaf and root tissues of Nisqually-1.

**Tables S5.** Primers used for qPCR and semi-quantitative RT-PCR analyses.

**Tables S1 to S4**, provided as a separate Excel file

**Supplemental Methods**

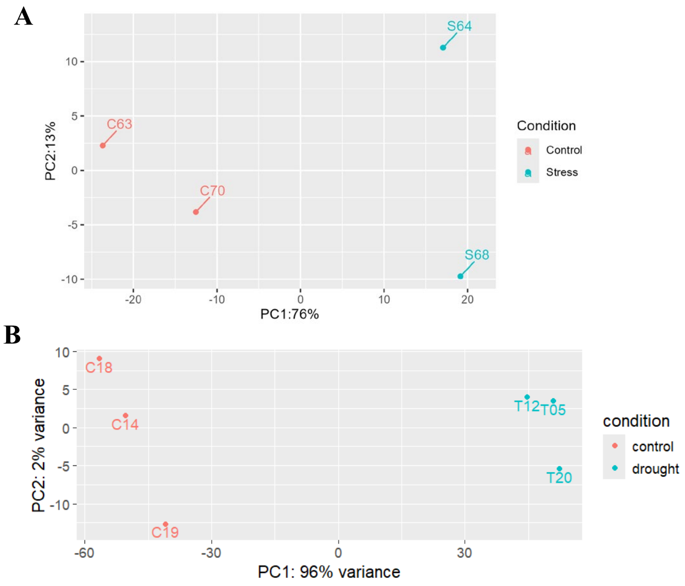


**Figure S1**. Principal component analysis (PCA) of gene expression in BCW (A) and 717 (B) under drought stress. Gene expression analysis was performed using DESeq2 with Illumina short-read sequencing data to confirm the drought effects in samples used for Nanopore DRS analysis.

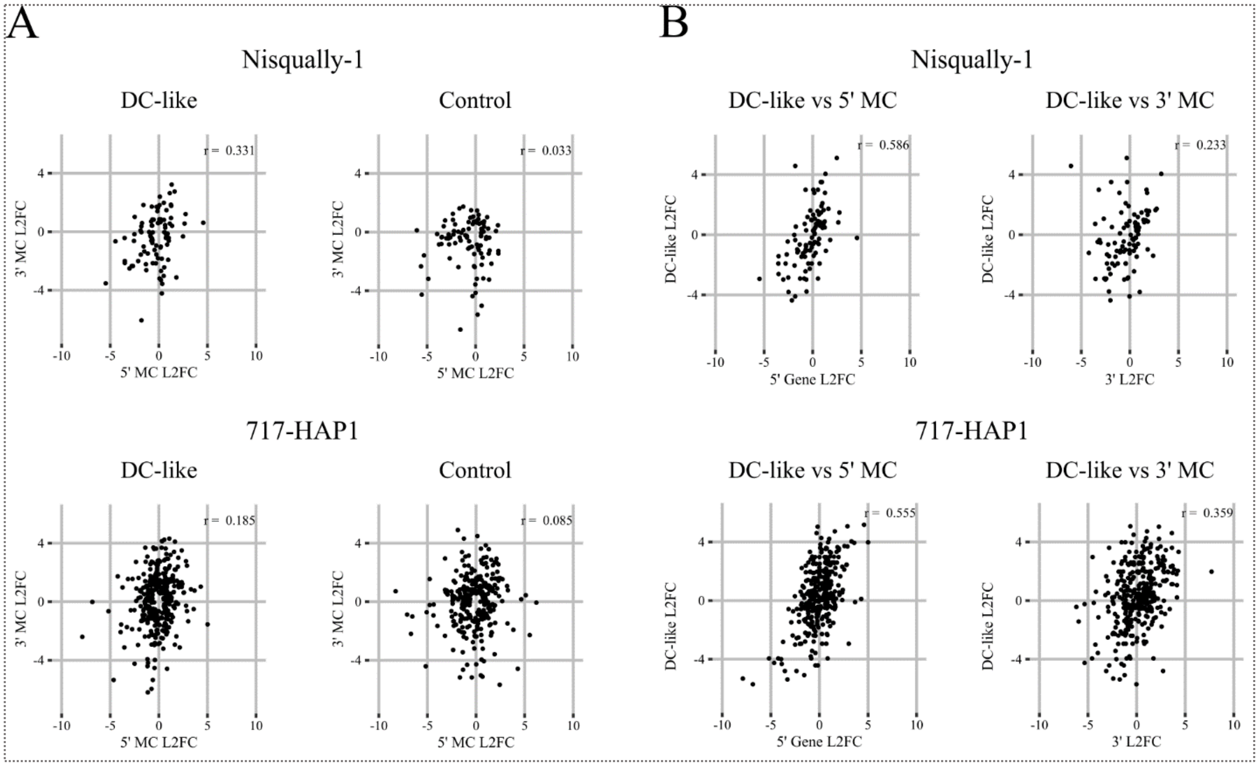


**Figure S2. Pearson correlation analysis of the expression of PC-like gene loci in Nisqually-1 mature leaf and hybrid 717 datasets using $\log_2$ fold changes (L2FC). A.** Pearson correlation analysis of the expression of monocistronic (MC) gene pairs at DC-like loci, compared with a control dataset of gene pairs in the same (sense) orientation that are not spanned by polycistronic-type RNAs. **B.** Pearson correlation analysis of the expression of PC-like RNAs with their corresponding 5′ monocistronic (MC) genes (left panel) and 3′ MC genes (right panel).

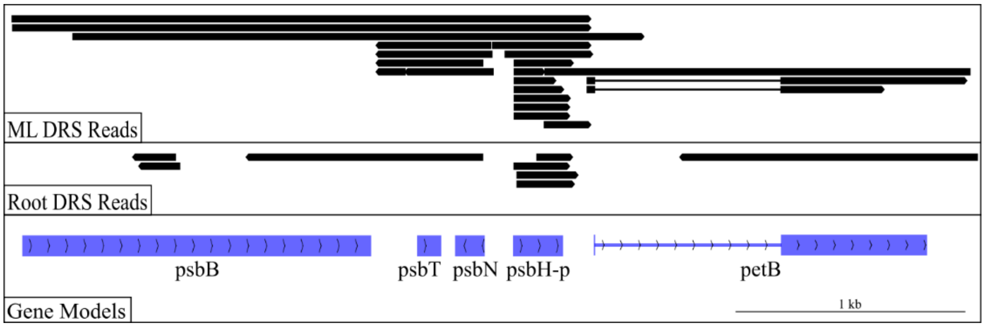


**Figure S3.** Detection of transcription of the chloroplast-encoded *psbH* gene using a poly(mA)-tailed total RNA DRS library (Yuan et al., 2025) from mature leaf (ML) and root tissue of Nisqually-1, demonstrated by DRS read alignments spanning the gene models within a polycistronic unit containing *psbH* in the chloroplast genome.

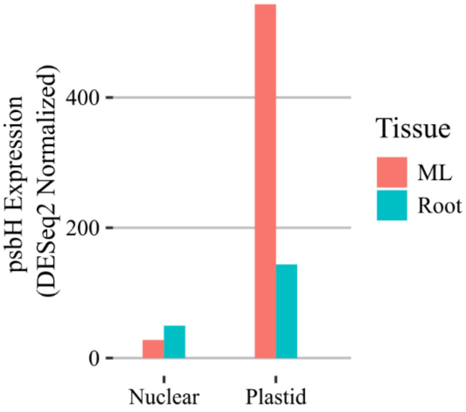


**Figure S4. Expression of nuclear NUPT *PsbH* and plastid *psbH* across tissues of Nisqually-1.** Read count matrices from featureCounts were generated for both nuclear-aligned reads and plastid-aligned reads and were further processed in R using DESeq2 to generate normalized data.

### Supplemental Table

**Table S5. Primers used for qPCR and semi-quantitative RT-PCR analysis**

| Primer | Sequence (5' -> 3') | Usage | Figures associated |
|---|---|---|---|
| PC13-qRT-F | ATCGGCATGATGGGTTGCCT | Realtime qPCR to quantify polycistronic RNAs spanning Potri.002G149800 and Potri.002G149900 | 3B |
| PC13-qRT-R | TTCTGATCCTGACACGCTCG | | |
| PC31-F | GGTTGATCCTAAAGGCCCGT | Realtime qPCR to quantify polycistronic RNA spanning Potri.016G136100 and Potri.016G136000 | 3D |
| PC31-R | GTCTGTCAGTGCCCCATTGC | | |
| T10_US_qRT_F | CTCTTTTCAGTGCCACCAAAAC | Realtime qPCR to quantify the DC loci Potri.009G111000-Potri.009G110901 | 1E |
| T10_US_qRT_R | CGCCAAAACTTTCCGCTGTT | | |
| T10_DS_qRT_F | TGTGCACTGGATTTACACGG | Realtime qPCR to quantify the DC loci Potri.009G110901-Potri.009G110800 | 1E |
| T10_DS_qRT_R | TCGTTCAACAGCTTTCCCAC | | |
| PC-7-FULL-F | AGAAGCAAGTTCAGTAATGGC | Semi-quantitative RTPCR to amplify tricistronic RNAs spanning Potri.011G142200, Potri.011G142300, and Potri.011G142501 | 4D |
| PC-8-Short-R | CGGAGACAGGAATGACCAA | | |
| Tar_1_H1SP | GTGCAGTCAAAAGGGAAAGATTTG | Realtime qPCR to quantify allele-specific expression of the dicistronic loci containing sugar-transporter and a plastid mobility related gene. The "Shared" primer was used with each allele-specific primer. | 2b |
| Tar_1_H2SP | GTGCAGTCAAAAGGGAAAGATTTC | | |
| Tar_1_Shared | AGTTACCAATGCAAGCAGCAATAG | | |
| UBQ-F | TGCGTGGAGGAATGCAAATC | Realtime qPCR of P. trichocarpa to amplify Potri.011G134200, a *UBQ* gene. Used as the reference gene for normalization in the ΔΔCt methods. | 2C, 3B, 3D, 4D |
| UBQ-R | CCTTTGTTGGTCAGGGGGTA | | |
| 18S rRNA-F | TCCTAGTAAGCGCGAGTCAT | Realtime qPCR of 717 to amplify nuclear 18S rRNA. Used as the reference gene for normalization in the ΔΔCt methods. | 2B |
| 18S rRNA-R | GAACACTTCACCGGACCATT | | |

### Supplemental Methods

#### Preparation of poly(mA) total RNA library for Nanopore DRS

To capture RNAs transcribed from chloroplasts and mitochondria for Nanopore DRS analysis, we employed a poly(mA) tailing-based Nanopore DRS method developed in our laboratory (Yuan et al. 2024), as organellar RNAs are generally not polyadenylated like nuclear RNAs.

For library preparation, 30 µg of DNase I-treated total RNA from *P. trichocarpa* was subjected to rRNA depletion using the riboPOOL kit for plants (siTOOLs Biotech, dp-K012-000031) according to the manufacturer's instructions. The resulting rRNA-depleted RNA was further cleaned and concentrated using the RNA Clean & Concentrator kit (Zymo, R1016).

Approximately 300 ng of rRNA-depleted RNA was used for the poly(mA) tailing reaction. RNA samples were first heated to 65 °C for 5 min to denature secondary structures, then subjected to a 20 µl reaction containing 600 units of yeast poly(A) polymerase (Thermo Fisher Scientific, 74225Z25KU), 1x yeast poly(A) polymerase reaction buffer, and 1 mM 2′-O-methyladenosine-5′-triphosphate (2′-OMeA; TriLink, N-1015-10). Reactions were incubated at 37 °C for 60 min and terminated by the addition of EDTA to a final concentration of 5 mM.

The tailed RNA was purified using RNAClean XP beads (Beckman Coulter, A63987), and eluted in 8 µl of water for subsequent library preparation. To ensure complete removal of residual 2′-OMeA, the eluted tailed RNA was further desalted using Performa spin columns (EdgeBio, 73328) according to the manufacturer's instructions.

Around 50 to 100 ng of poly(mA)-tailed RNA is used for Nanopore DRS library construction following the manufacture's instructions. The resulted direct RNA libraries are sequenced on PromethION RNA flow cells (Oxford Nanopore Technologies, FLO-PRO004RA) using a

PromethION 2 Solo (P2 Solo) device connected to a GridION Mk1 as a compute resource and operated with standard MinKNOW software (version 24.11.8).
